# Deciphering how naturally occurring sequence features impact the phase behaviors of disordered prion-like domains

**DOI:** 10.1101/2021.01.01.425046

**Authors:** Anne Bremer, Mina Farag, Wade M. Borcherds, Ivan Peran, Erik W. Martin, Rohit V. Pappu, Tanja Mittag

## Abstract

Phase separation of intrinsically disordered prion-like low-complexity domains (PLCDs) derived from RNA-binding proteins enable the formation of biomolecular condensates in cells. PLCDs have distinct amino acid compositions, and here we decipher the physicochemical impact of conserved compositional biases on the driving forces for phase separation. We find that tyrosine residues make for stronger drivers of phase separation than phenylalanine. Depending on their sequence contexts, arginine residues enhance or weaken phase separation, whereas lysine residues weaken cohesive interactions within PLCDs. Increased net charge per residue (NCPR) weakens the driving forces for phase separation of PLCDs and this effect can be modeled quantitatively. The effects of NCPR also weaken known correlations between the dimensions of single chains in dilute solution and the driving forces for phase separation. We build on experimental data to develop a coarse-grained model for accurate simulations of phase separation that yield novel insights regarding PLCD phase behavior.

---

Biomolecular condensates enable spatial and temporal organization of biochemical reactions that control key cellular processes such as division ^1^, signaling ^2^, and transcriptional regulation ^3^. Examples of condensates include ribonucleoprotein condensates such as stress granules that sequester naked, unfolded RNA molecules when stress causes the termination of translation and polysomes disassemble ^4^. Stress granule formation and dissolution is regulated by reversible phase transitions that are regulated by a network of heterotypic protein-RNA interactions and homotypic protein-protein interactions ^5, 6, 7^.

Many of the protein components of stress granules are RNA-binding proteins (RBPs) that feature tandem RNA-binding domains and intrinsically disordered prion-like low complexity domains (PLCDs) ^8, 9^. Mutations within PLCDs can cause neurodegenerative diseases such as Amyotrophic Lateral Sclerosis (ALS) ^10, 11^, and genetic translocation events that result in the expression of PLCD fusion proteins can cause multiple cancers ^12, 13^. Mutations appear to impact the phase behavior of PLCDs and there is growing consensus that aberrant phase transitions underlie disease processes ^14, 15^. Proteins that are genetically linked to ALS include RBPs such as TIA-1, TAF15, FUS, TDP-43, EWSR1, hnRNPA1, hnRNPA2, and hnRNPD/L. Mutations associated with ALS tend to be located within PLCDs that can phase separate as autonomous units ^14, 16^. *In vitro* investigations of PLCD phase behavior reveal that disease-related mutations can alter the driving forces for phase separation, change material properties of condensates, or enable the conversion from liquid-like condensates to fibrillar solids ^11, 14, 15^. It is therefore critical to understand how the driving forces for functional and aberrant phase separation are encoded in PLCDs.

The *stickers-and-spacers* framework has proven to be useful for describing the normal and aberrant phase behaviors of PLCDs and other multivalent protein and RNA molecules that encompass disordered regions and folded domains ^16, 17, 18, 19^. In this framework, stickers in disordered regions are short linear motifs or individual residues that make reversible, non-covalent crosslinks with one another. Depending on the identities of the stickers, the physical crosslinks involve a combination of ionic, multipolar, hydrophobic, cation-pi, and pi-pi interactions ^20, 21, 22, 23, 24^. Accordingly, physical crosslinks have characteristic length and energy scales ^19, 25^. Above a concentration threshold known as the percolation threshold, physical crosslinks can lead to the formation of a system-spanning network ^18, 19, 26^. Spacers provide scaffolds for stickers, and the effective solvation volumes of spacers determine whether percolation is aided by phase separation or if percolation arises without phase separation ^19, 27, 28^. Phase separation-aided percolation transitions give rise to coexisting dilute and dense phases, where the latter are condensates that are best described as viscoelastic network fluids ^18, 28, 29^.

In a recent study, directed toward the PLCD from isoform A of human hnRNPA1, referred to hereafter as the WT A1-LCD or A1-LCD, aromatic residues were shown to be the primary stickers that form reversible non-covalent crosslinks ^17^. Stickers are uniformly distributed along the linear sequence and this leads to a dilution of the effects of inter-sticker crosslinks by spacers. The valence (number) of stickers and their patterning along the linear sequence determine the phase behavior of archetypal PLCDs such as the A1-LCD. The interplay among sticker-sticker, sticker-spacer, and spacer-spacer interactions leads to percolated condensates with liquid-like properties that dissolve above an upper critical solution temperature (UCST) ^17^.

The binary classification of PLCD residues as aromatic stickers and non-aromatic spacers suggests that aromatic residues such as Tyr and Phe are interoperable with one another. It also suggests that non-aromatic residues are unlikely to contribute as stickers, and that all non-aromatic residues might be interoperable with one another as spacers. If these implications are true, then given sufficient evolutionary drift ^30^ and no selection pressure on the identities of non-aromatic residues, the amino acid compositions of PLCDs from homologous proteins should feature a conservation of the valence and patterning of aromatic stickers and a lack of conservation of any other sequence features. To assess if this expectation is valid, we quantified the amino acid composition of A1-LCD (Fig. 1a) and analyzed the extent to which compositions are conserved across 848 PLCDs drawn from orthologs and paralogs (referred to hereafter as homologs) within the hnRNPA1 family (Fig. 1b). Sequences for PLCDs of hnRNPA1 were drawn from proteomes of unrelated organisms, and include species that diverged ~750 M years ago ^30^. We reduced each of the 848 PLCD sequences to a 20×1 vector that defines its amino acid composition and calculated the similarities of amino acid compositions between all pairs of PLCDs from homologs. This is quantified by computing the cosine of angles between every pair of vectors. Most of the cosine values are close to one (Fig. 1b) implying that compositional biases are similar across PLCDs from homologous proteins. This suggests that there are conserved preferences for non-aromatic residues. It also indicates that binary classifications into aromatic stickers and non-aromatic spacers might be too coarse-grained for describing the molecular grammar underlying the driving forces for phase separation of PLCDs.

**Fig. 1:**
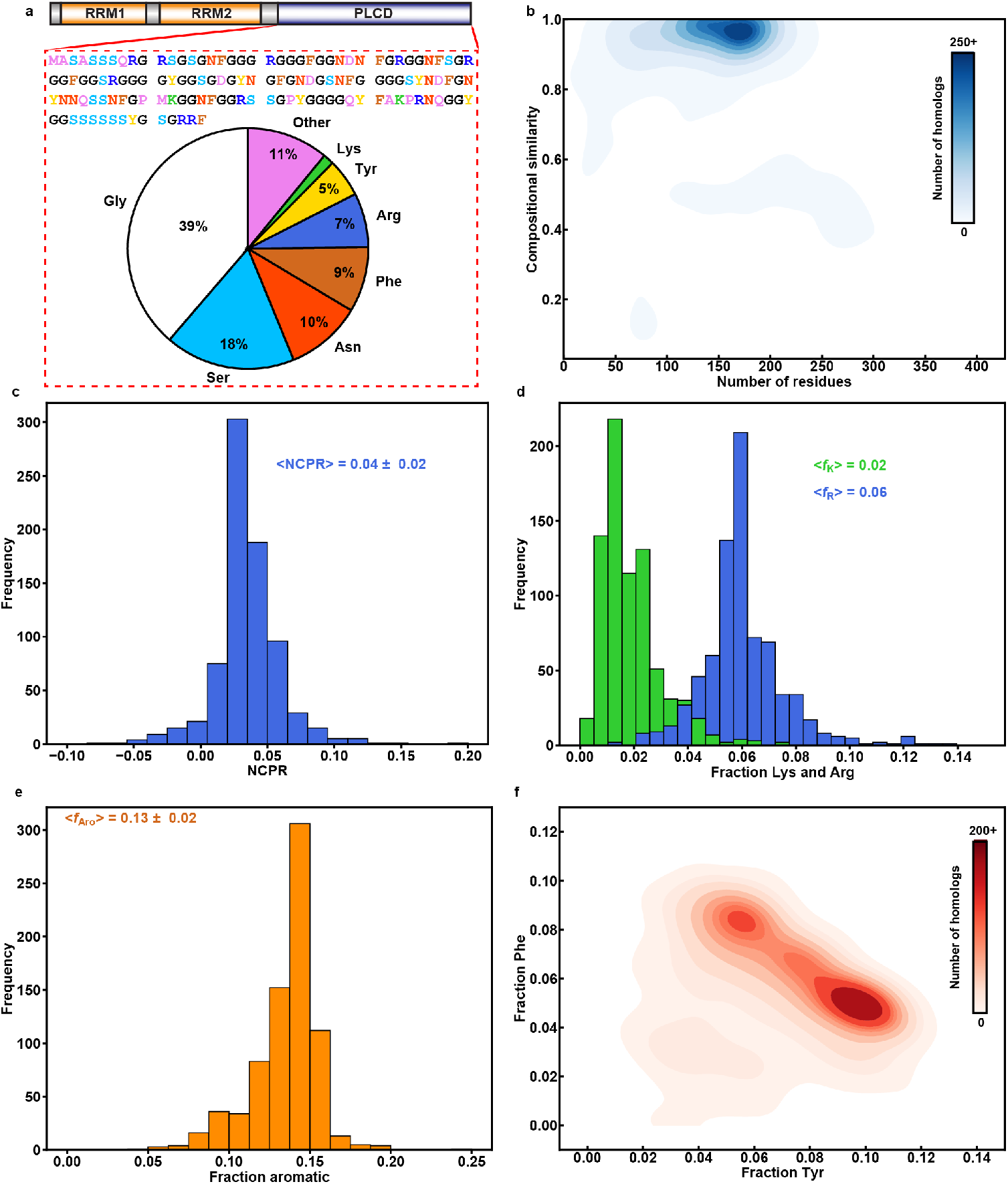
Compositional analysis of PLCDs from homologs of hnRNPA1. **(a)** Amino acid sequence and compositional statistics for the PLCD from isoform a of hnRNPA1 – referred to in the text as A1-LCD. **(b)** 2D histogram quantifying the joint distribution of lengths of PLCDs derived from homologs of hnRNPA1 and the compositional similarities, in terms of cosine values, to A1-LCD. **(c)** Distribution of NCPR values; **(d)** Distributions of fractions of Arg and Lys residues; **(e)** Distribution of fractions of aromatic residues; Numerical values are mean values of the respective distribution ± the standard deviations. **(f)** 2D histogram quantifying covariations in fractions of Tyr versus Phe residues across PLCDs.

To motivate a design-based approach for understanding the sequence-to-phase-behavior relationships of PLCDs, we extracted features that characterize the compositions of PLCDs across homologs. Key parameters include the net charge per residue (NCPR) (Fig. 1c), the fractions of Arg versus Lys (Fig. 1d), the fraction of aromatic residues (Fig. 1e), and the covariations between Tyr and Phe contents (Fig. 1f). These analyses were motivated by results that highlight the role of Arg as a sticker ^16^, the relative weakness of Lys as a sticker ^16, 22, 23, 31, 32^, the contributions of charged residues that destabilize phase separation either as spacers that promote solvation ^27, 28^ or via electrostatic repulsions ^33^, and the suggestion that Tyr is likely to be a stronger sticker than Phe ^16, 34^.

Analysis of the compositional biases across PLCDs from homologs of hnRNPA1 shows that NCPR values are narrowly distributed around a mean value of +0.04 (Fig. 1c). There is a clear preference for Arg over Lys, with the Arg and Lys contents being narrowly distributed around mean values of 0.06 and 0.02, respectively (Fig. 1d). The fraction of aromatic residues is narrowly distributed around a mean value of 0.13 (Fig. 1e). Further analysis shows that the fractions of Phe and Tyr residues vary considerably across A1-LCD sequences from homologs. Specifically, there is a clear negative correlation between the fractions of Phe and Tyr residues across the 848 sequences indicating a covariation between fractions of Phe and Tyr (Fig. 1f,g). Our results point to a complex compositional landscape, with clear biases for heterogeneous distributions of non-aromatic residues. While our bioinformatics analysis uncovers distinct compositional preferences, it does not provide insights into the physical basis for these trends. Here, we used a combination of biophysical investigations, machine learning-aided ^35^ coarse-grained simulations ^18^, and theoretical analysis that collectively yield a physicochemical rationalization for the conserved biases and covarying trends across A1-LCD homologs. We uncover molecular explanations for previous observations and for previously unappreciated context dependencies of compositional effects. Building on rules that emerge from our systematic efforts, we propose, based on additional evolutionary analysis of PLCDs from other FET family proteins, that the rules we have extracted from this work are transferrable across a wide range of PLCDs.

In what follows, we present results from systematic investigations that uncover the physical significance for NCPR being narrowly dispersed about a mean value of +0.04, the rationale for there being a clear preference for Arg over Lys residues, and the consequences of varying Phe versus Tyr contents while keeping the fraction of aromatic residues and hence the valence of aromatic stickers narrowly dispersed. Additionally, PLCDs are flexible, finite-sized heteropolymers, and yet they have been shown to behave in a manner that is similar to effective homopolymers ^17^, whereby the determinants for the collapse of each chain in dilute solutions are the same as those for driving phase separation ^36^. This “strong coupling” between the determinants of conformational and phase equilibria of PLCDs has been attributed to the uniform distribution of aromatic stickers along the linear sequence ^17, 37^. The sequence titrations that we deploy in this work allow us to quantify the impact of changes to sticker versus spacer grammars on the coupling between single-chain dimensions and the driving forces for phase separation. Finally, we use our computational model, parameterized to reproduce the totality of experimental observations, to assess the differences and similarities of single chain conformations in the dense versus dilute phases and uncover the interplay between overlap concentrations that are governed by conformational fluctuations and the cohesive effects of inter-sticker interactions ^38^.

## Results

### Design of sequence variants of A1-LCD

Motivated by our analyses of compositional biases and covariation trends across homologs, we generated 28 distinct sequence variants of the A1-LCD (Fig. 2a, Table S1). All sequence variants have the same length. The designations used in Fig. 2a reflect the numbers of aromatic / charged residues that were added or removed. Additions or deletions of aromatic / charged residues were achieved either by replacing Gly / Ser spacers with the residues of interest or substituting deleted residues with Gly / Ser residues. These additions and deletions preserve the overall balance of Gly and Ser, which are the most abundant residues in A1-LCD (Fig. 1a) and are thought to be spacers as opposed to stickers ^16^.

**Fig. 2:**
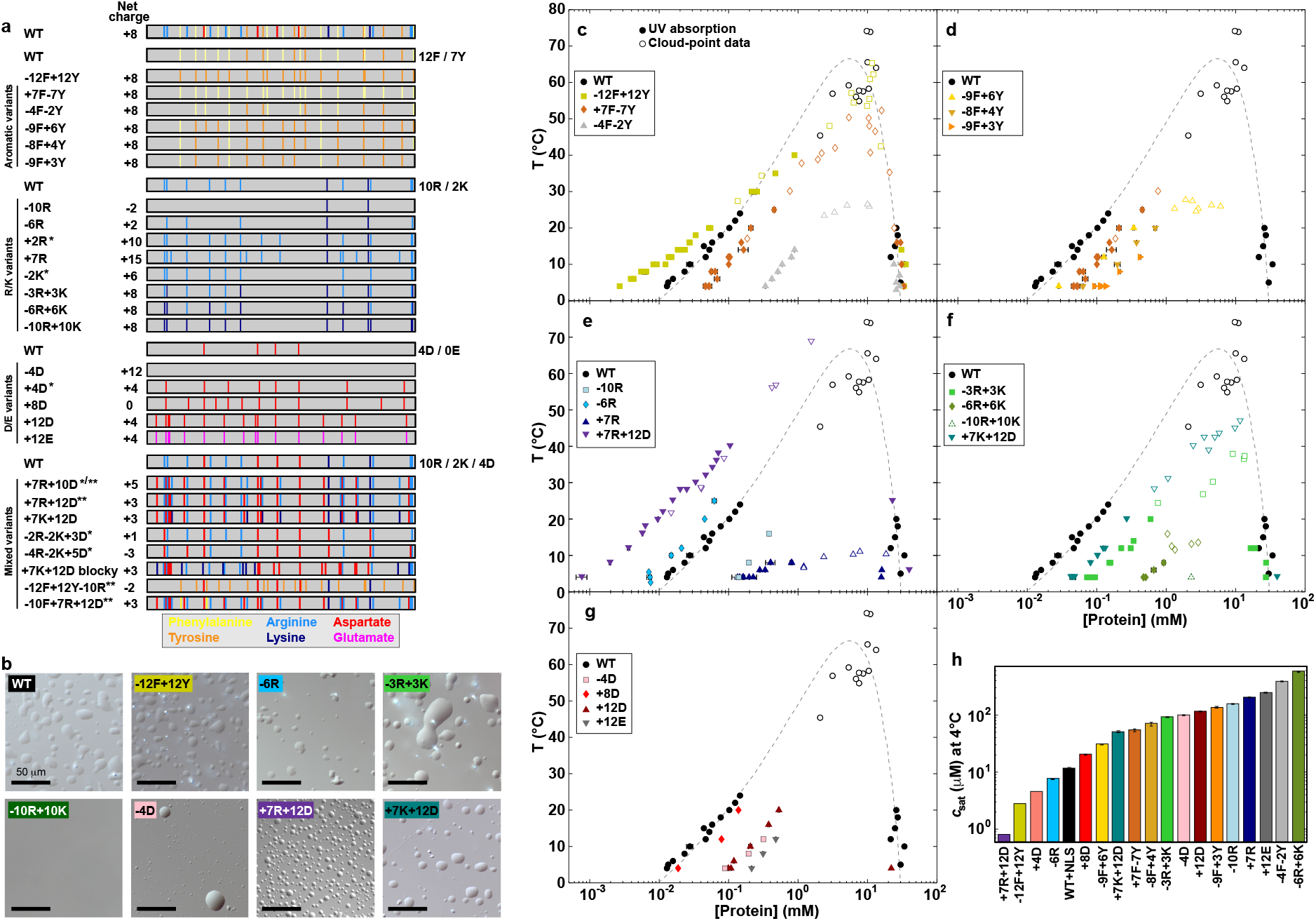
Comparative assessments of sequence-binodal relationships for designed variants of A1-LCD. **(a)** A1-LCD variants that titrate aromatic and charged residues. Numbers next to the schematic for WT indicate the number of residues of a certain type. Vertical bars in the schematics indicate the position of residue types namely, Phe (yellow), Tyr (orange), Arg (light blue), Lys (dark blue), Asp (red), and Glu (magenta). Asterisks indicate A1-LCD variants used to test the validity of the analytical model in Fig. 3 (*) and the coarse-grained model in Fig. 6 (**). **(b)** Differential interference contrast (DIC) images of select A1-LCD variants showing that they form dense liquid droplets. Scale bar represents 50 μm. Solution conditions were 20 mM HEPES, 150 mM NaCl, pH 7.0. **(c-g)** Binodals of A1-LCD variants as a function of temperature in 20 mM HEPES, 150 mM NaCl, pH 7.0. Binodals of A1-LCD for aromatic (c, d), Arg (e), Arg/Lys (f), Asp/Glu (g), and mixed variants where the number of positively and negatively charged residues is altered (e,f). Binodals were determined using two different types of experiments *viz*., centrifugation followed by UV absorbance measurements (filled symbols) and cloud point measurements (open symbols). The dashed line in each of the panels (c)-(g) is a fit of Flory-Huggins theory to the experimental data for the WT A1-LCD. Data for -4F-2Y are those of Martin et al. **(h)** Summary of variantspecific saturation concentrations measured at 4°C. Error bars indicate the standard error of replicate measurements.

We expressed and purified each of the 28 variants (Fig. S1) and characterized their temperature-dependent phase behaviors *in vitro*. All variants (except for -10R+10K) form liquid-like condensates giving rise to dense, liquid-like droplets at pH 7.0 and 4°C in the presence of 150 mM NaCl. Differential interference contrast micrographs are shown in Fig. 2b for a subset of the variants. The variants can be grouped into five distinct classes that allow us to assess the impact of specific sequence features on the driving forces for phase separation as measured by coexistence curves. With the aromatic variants (Fig. 2a) we query the impact of variations to Tyr versus Phe contents. With the R-variants (-10R, -6R, and +7R) we assess whether Arg residues contribute as auxiliary stickers in sequences where aromatic residues act as the primary stickers. The R/K variants (3R+3K, 6R+6K, and -10R+10K) help us query the impact of systematically replacing Arg with Lys. With D/E variants (-4D, +8D, +12D, +12E), we assess the contributions of changes to the effective solvation volumes of spacer residues. Finally, the mixed variants (+7R+12D and +7K+12D) allow us to interrogate the effect of increasing the fraction of charged residues by jointly increasing the fraction of acidic (Asp) and basic (Lys or Arg) residues.

### Impact of Tyr versus Phe on the driving forces for phase separation

We used sedimentation assays as described previously ^17, 39^ to measure coexisting dilute and dense phase concentrations as a function of temperature. Cloud point measurements were used to identify the locations of critical points ^40^. Fig. 2c shows the measured coexistence curves and cloud points for the aromatic variants. The WT A1-LCD has 12 Phe and 7 Tyr residues distributed uniformly along the sequence (Fig. 2a). In the -12F+12Y variant, all Phe residues of the WT are replaced with Tyr residues. This increases the width of the two-phase regime, although the apparent critical point and the dense phase concentration stay roughly the same as those of the WT. Widening of the two-phase regime is achieved by shifting the low concentration arm (left arm) of the binodal to lower concentrations. This implies that substituting Phe with Tyr residues enhances the driving forces for phase separation without causing changes to the density within condensates. The variant +7F-7Y replaces all Tyr residues with Phe. In contrast to the -12F+12Y variant, the width of the two-phase regime narrows for +7F-7Y. We measured coexistence curves for -9F+6Y, -8F+4Y, and -9F+3Y, which contain Phe and Tyr residues but shift their ratios. The data show that the overall phase behavior is determined by a combination of the total number of aromatic stickers and the asymmetry between Tyr versus Phe residues (Fig. 2d).

### Comparative assessments of the phase behaviors of different charge variants

The favorable free energies of solvation of charged residues should increase the effective solvation volumes of spacers and destabilize phase separation ^27, 28^. In addition, Lys lacks a quadrupole moment and is likely to behave differently from Arg ^41, 42^. We performed a series of measurements to obtain comparative assessments of how replacements or additions of different types of charges contribute to the phase behaviors of charge variants of A1-LCD. First, we assessed the phase behavior of the R-variants -10R, -6R, and +7R. The coexistence curve of -10R, which removes all 10 Arg residues from the sequence, changed considerably when compared to the A1-LCD, weakening the driving force for phase separation in line with previous work that has identified Arg residues as drivers of phase separation ^16, 23^. However, to our surprise the -6R variant shows a slight increase in the width of the two-phase regime compared to WT, whereas phase separation is significantly destabilized for +7R (Fig. 2e). Together, these data suggest that Arg residues contribute as auxiliary stickers within PLCDs, but these contributions are context-dependent resulting in non-trivial relationships between Arg content and the driving forces for phase separation. Next, we assessed the effects of substituting Arg with Lys residues in variants -3R+3K, -6R+6K, and -10R+10K. These substitutions clearly weaken the driving forces for phase separation (Fig. 2f). In fact, replacement of all Arg residues with Lys (-10R+10K) resulted in the abrogation of phase separation in the concentration range up to 2.3 mM.

To assess the impact of acidic residues on phase separation, we added 8 or 12 Asp residues to generate variants +8D and +12D, or 12 Glu residues to generate +12E, to the 4 Asp residues in the WT sequence. Increasing the Asp or Glu content beyond that of the WT sequence leads to substantial weakening of the driving forces for phase separation. This is consistent with the hypothesis that increasing the charge content, without compensatory electrostatic attractions, destabilizes phase separation through the increased effective solvation volumes of and electrostatic repulsion among spacer residues (Fig. 2g). Interestingly, substitution of the four native Asp residues with Gly / Ser, as in variant - 4D, also weakened the driving forces for phase separation. This points to a buffering role of the native Asp residues because replacement of these groups increases the uncompensated positive charge in the system.

To test the hypothesis that electrostatic attractions offset the destabilizing effects of high effective solvation volumes, we measured coexistence curves for variants +7R+12D and +7K+12D. Indeed, the destabilizing effects seen in +7R and +12D are offset and overcome in the constructs that are close to being electroneutral (Fig. 2e,f). This effect also prevails for the +7K+12D variant, although the differences between Arg and Lys still persist. Comparison of the coexistence curves for +7R+12D and +7K+12D shows that a combination of near electroneutrality and the presence of context dependent auxiliary sticker effects of Arg contribute to the enhancement of phase separation in +7R+12D when compared to the A1 - LCD. In summary, the sequence variation in this set of A1-LCD variants modulated the saturation concentration by three orders of magnitude (Fig. 2h), and we sought a framework to understand the molecular underpinnings of these sequence-dependent changes to the driving forces for phase separation.

### Application of a mean-field model explains the data for charge variants

We hypothesized that electroneutrality might be important for enhancing the driving forces for phase separation. According to this hypothesis, the WT sequence with its 4 Asp, 10 Arg, 2 Lys, and a net charge of +8 should have an unfavorable balance of charged residues. We assessed the impact of net charge by plotting the measured log_10_(*c*_sat_) values against NCPR for a series of variants with identical aromatic stickers but changes to charged residues (Fig. 3a). Here, *c*_sat_ is the coexisting dilute phase concentration known as the saturation concentration. We performed this analysis using data collected at 4°C. The plot of log_10_(*c*_sat_) versus NCPR at 4°C has a putative V-shaped profile (Fig. 3a) indicating the possibility that *c*_sat_ is at its lowest value for electroneutral systems (NCPR ≈ 0). However, differences between Arg and Lys variants lead to significant deviations from the putative V-shaped profile (Fig. S2a). Specifically, for equivalent NCPR values, we find that increasing the number of Arg residues lowers *c*_sat_ whereas decreasing the number of Arg residues increases *c*_sat_. In contrast, increasing the number of Lys residues increases *c*_sat_ (+7K+12D, -3R+3K, and -6R+6K) to extents that cannot be explained by deviations from electroneutrality.

**Fig. 3:**
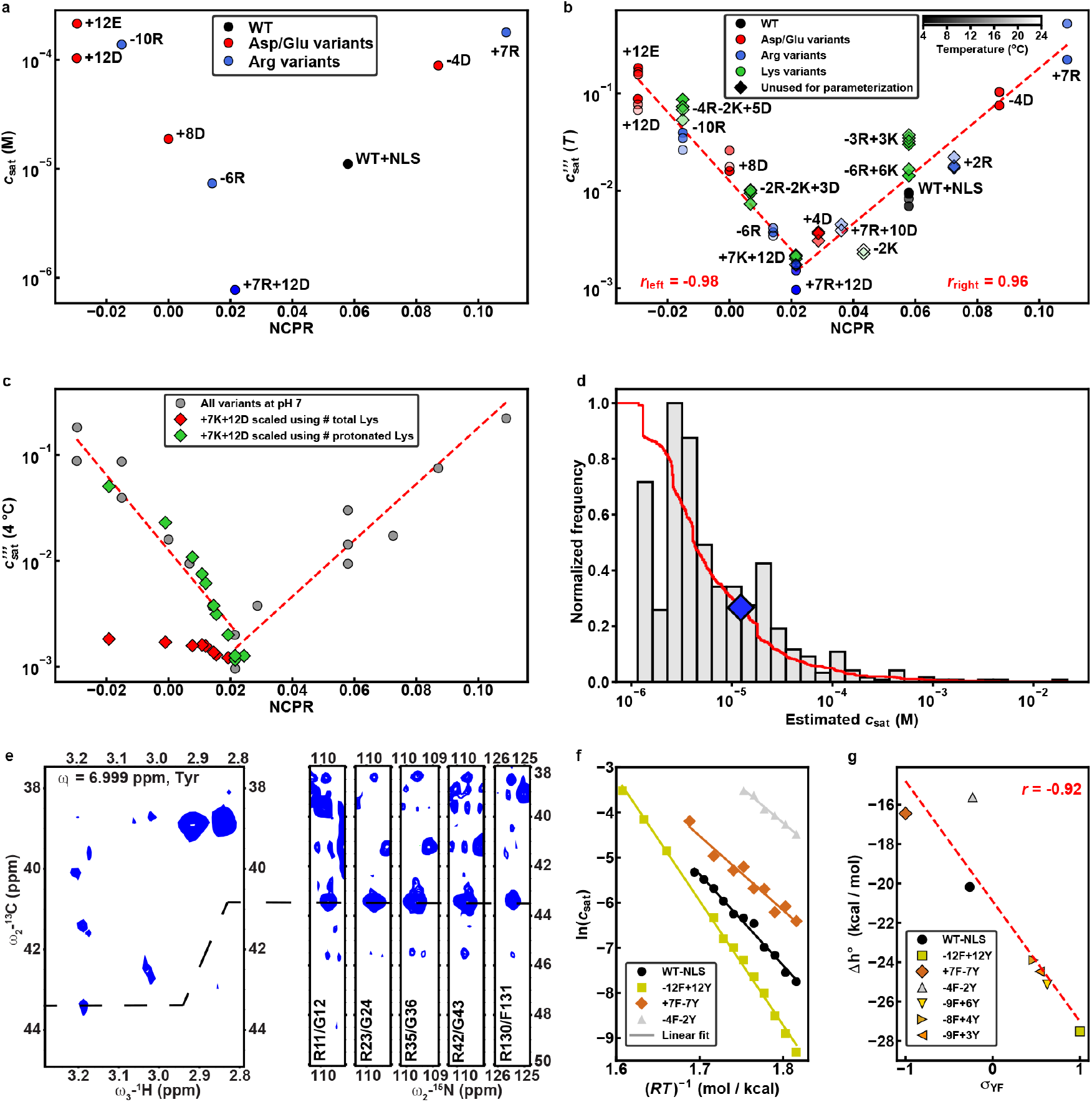
NCPR impacts A1-LCD phase separation. **(a)** log_10_(*c*_sat_) from data measured at 4°C, plotted against NCPR for a subset of the sequence variants. **(b)** Rescaled *c*_sat_ i.e., 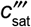, from data measured at multiple temperatures, plotted against NCPR. The validity of the model is tested with variants (diamond markers) that were not used in the parameterization. Left and right red dashed lines are linear fits to the data corresponding to variants with NCPR less than or equal to that of +7R+12D and greater than or equal to that of +7R+12D, respectively. (**c**) Measured *c*_sat_ values for +7K+12D were rescaled using the equation for 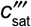. Here, we compare the 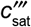 values when we account for all nine Lys residues (red diamonds) or only the number of protonated Lys residues (green diamonds). The latter conform to the master curve, whereas the former deviate significantly from the master curve. (**d**) Histogram of calculated *c*_sat_ values for 640 PLCDs from the set of 848 sequences derived from hnRNPA1 homologs. These *c*_sat_ values were calculated using the mean-field model that yields 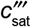 in panel (b). The red curve is the cumulative distribution function that quantifies the probability that PLCDs have specific *c*_sat_ values. For example, the probability that a PLCD has a *c*_sat_ value that is less than or equal to that of the WT A1-LCD (blue diamond) is ≈ 0.78. **(e)** Tyr sidechain plane of ^13^ C-resolved ^1^H^aromatic_1^H^aliphatic^ NOESY spectrum (left) shows NOEs between sidechain protons of Tyr and H^δ^ protons of Arg (right). Degenerate Arg H^δ^ proton resonances are shown as strips of (H)CC(CO)NH spectrum (right; see Fig. S6). **(f)** Results from van’t Hoff analysis of select A1-LCD aromatic variants. **(g)** Inferred values of Δh° plotted against σ_YF_ for all A1-LCD aromatic variants used in this study. Red dashed line is the linear fit to the data.

We reasoned that Arg residues are context-dependent auxiliary stickers, whereas Lys residues weaken attractive interactions ^26, 43^. To test this hypothesis, we used a generalized mean-field stickersand-spacers model ^26^, and rescaled the *c*_sat_ values at 4°C using 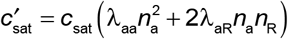. In this formalism, we account for the contribution of Arg residues as auxiliary stickers by fixing λaa = 1, and use regression analysis to obtain λ_aR_, which is 1.69. Higher l-values with a positive sign imply less favorable contributions from aromatic-Arg (aR) interactions compared to inter-aromatic (aa) interactions; *n*_a_ and *n*_R_ quantify the numbers of aromatic and Arg residues, respectively. To account for residuals that persist in the plot of *c*’_at_ against NCPR (Fig. S2b) we accounted for the effects of Lys residues in a further rescaling 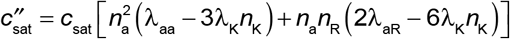. Regression analysis yields λ_K_ = 0.0479, and the minus sign for terms involving λ_K_ quantifies the unfavorable contributions of Lys; here, *n*_K_ is the number of Lys residues. With this rescaling, all data points at 4°C can be collapsed onto two arms of a single V-shaped plot (Fig. S2c). Finally, we account for the changes to *c*_sat_ with temperature to arrive at 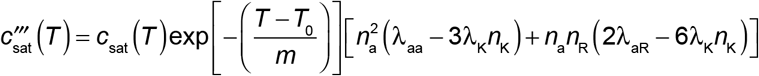 that yields a master Curve for 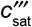 versus NCPR (Fig. S2d, 3b) where we set *T*_0_ = 277 K and obtain the optimal value of *m* = 8.26 K.

To test predictions made by the master V-shaped plot, we generated six additional variants (marked with asterisks in Fig. 2a). These variants fill in the gaps on the V-shaped master curve. Binodals for these new variants are shown in Fig. S3. Importantly, without any further reparameterization, we find that the measured *c*_sat_ values for the new variants are readily rescaled to coincide with predictions from the V-shaped master curve (Fig. 3b). The V-shape of the master curve results from the destabilizing effects of increased net charge. However, we find that the minimum in the V-shaped profile is located at a positive value of NCPR as opposed to being at zero. All NCPR values were calculated by assuming unshifted pK_a_ values for ionizable groups. To test for the possibility that the theoretical and actual NCPR values might be different due to shifted pK_a_ values of ionizable residues, we measured the pH dependence of phase separation for the +7K+12D variant. All data points collapse reasonably well onto the V-shaped master plot when we rescaled the measured *c*_sat_ values by estimating the expected numbers of protonated Lys residues as the entities that weaken inter-sticker interactions (Fig. S4, Fig. 3c). These observations suggest that the true minimum of the V-shaped plot does indeed coincide with positive NCPR values.

We used the mean-field model, specifically the expression for 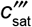 and the values obtained for the λcoefficients to estimate *c*_sat_ at 4°C for PLCD sequences derived from homologs of hnRNPA1 (Fig. 3d). We restricted the analysis to PLCDs that were 100-200 residues long in order to ensure that changes to chain length do not confound the application of our mean-field model. The results in Fig. 3d show that our model predicts values for *c*_sat_ that span up to three orders of magnitude, although most of the PLCDs are predicted to have *c*_sat_ values that are in the low micromolar range. Specifically, from the cumulative distribution function, we find that the probability of finding PLCDs with *c*_sat_ values that are less than or equal to those of WT A1-LCD is ≈0.78.

A salient prediction from our model is that Arg residues are auxiliary stickers that interact with aromatic stickers. Inter-sticker interactions can be detectable as nuclear Overhauser enhancements (NOEs) using nuclear magnetic resonance (NMR) spectroscopy ^17^. NOEs are the result of magnetization transfer between nuclei through space and imply that physical distances between the nuclei are, at least transiently, within 5 Å of one another. Based on the established equivalence between cohesive interactions within single-chains and the driving forces for multi-chain phase behavior of A1-LCD variants ^17^, we assessed the presence of Arg-Phe/Tyr NOEs in NMR spectra in the dilute phase. A temperature of 40°C enabled a dilute phase concentration of 800 μM A1-LCD in the NMR sample, which was high enough to observe NOEs between aromatic sidechain protons of Phe / Tyr and arginine H^δ^ protons (Fig. 3e, Fig. S5). This supports the prediction that Arg residues interact with Phe / Tyr acting as auxiliary stickers within PLCDs.

NMR spectra for the +7K+12D variant showed large-scale changes in resonance frequencies compared to the A1-LCD (Fig. S6, S7). Additionally, *R*_2_ relaxation rates, which are sensitive to differences in local dynamics due to intramolecular interactions, showed clusters of enhanced rates in similar sequence positions as the WT, mainly pointing to aromatic residues as stickers (Fig. S7). These data support our prediction that Lys and Asp residues do not act as stickers. Instead, they modulate the driving forces for phase separation through a combination of increased effective solvation volumes, electrostatic repulsions, and weakening attractive interactions among primary and auxiliary stickers.

### Extracting thermodynamic parameters from the temperature dependence of *c*_sat_

Our measurements indicate that Tyr is an intrinsically stronger sticker than Phe ^16^. We used a van’t Hoff analysis method ^44^ to extract the apparent enthalpies (slopes) and entropies (intercepts) from plots of ln(*c*_sat_) versus (*RT*)^-1^ (see details in the Methods). The inferred parameters quantify the changes in standard state molar enthalpies (Δh°) and molar entropies (Δs°) associated with the transfer of PLCD molecules from the dilute to the dense phase (Fig. 3f, Fig. S8). Replacing all Phe residues with Tyr (-12F+12Y) resulted in larger negative (favorable) Δh° values compared to the A1-LCD. Conversely, replacing all Tyr residues with Phe (+7F-7Y) reduced the enthalpy change associated with phase separation. Shifting the balance between Tyr and Phe resulted in intermediate Δh° and Δs° values with a linear dependence on the Tyr versus Phe contents (Fig. 3g, Fig. S8). In this analysis, we use an asymmetry parameter defined as: 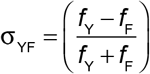 where *f*_Y_ and *f*_F_ are the fractions of Tyr and Phe, respectively. Note that σ_YF_ takes values of -1 for *f*_Y_ = 0, 0 for *f*_F_ = *f*_Y_, and +1 for *f*_F_ = 0.

We also performed van’t Hoff analyses on all variants using their measured temperaturedependent *c*_sat_ values (Fig. S8). We included an additional variant designated as +7K+12D blocky (Fig. 2a). In this variant, the oppositely charged residues are segregated into blocks along the linear sequence, while the positions of the aromatic residues are kept fixed. We used the data in Fig. S8 to assess the effects of Tyr/Phe-Arg interactions versus inter-aromatic interactions. We find that the Δh° values for - 12F+12Y and -12F+12Y-10R (Fig. S8e) are essentially identical to one another and have the largest magnitudes. This suggests that Tyr-Arg interactions contribute weakly when compared to Tyr-Tyr interactions. Interestingly, and in striking contrast, the standard-state enthalpy decreases by ~ 5 kcal/mol when we compare the WT A1-LCD to the -10R variant. This suggests that the Tyr/Phe-Arg interactions contribute significantly to the driving force for phase separation. Importantly, these results highlight the context dependence of Arg as an important versus irrelevant auxiliary sticker based on the Tyr versus Phe content within PLCDs.

Finally, the entropy changes (Δs°) associated with the transfer of PLCD molecules from the dilute to dense phase are always unfavorable i.e., values of -Δs°/R are positive in Fig. S8f. This is a signature of UCST behavior for the WT and all designed variants. The values of Δs° combine contributions from the gain / loss of solvent entropy, conformational entropy, the entropy of mixing, and the overall translational and rotational entropies. The results highlight the interplay between enthalpy and entropy whereby variants defined by large, favorable changes in enthalpy are opposed by higher negentropic penalties (Fig. S8e,f). This gives rise to a loss of two-phase behavior above the UCST and partly determines the variant-specific critical temperature values.

### Assessing the correlation between single-chain dimensions and *c*_sat_

Previous work showed that increasing the number of aromatic residues while maintaining the uniform linear distribution of these stickers leads to increased chain contraction in the dilute phase and lowering of *c*_sat_. The converse is true when the number of aromatic stickers is reduced ^17^. To zeroth-order, sticker valence, providing the patterning of stickers is uniform, appears to have equivalent effects on the cohesive interactions that control chain dimensions in the dilute phase and the driving forces for phase separation as measured by the values of *c*_sat_ and the widths of two-phase regimes. We asked if the correlation between single chain dimensions and *c*_sat_ is preserved across the wide range of variants studied here. These assessments test the robustness of the strong coupling between the driving forces for single-chain contraction and for phase separation of disordered proteins with sticker-and-spacer architectures.

We collected small-angle X-ray scattering (SAXS) data for dilute solutions of most of the variants. We leveraged size exclusion chromatography (SEC)-coupled SAXS to remove aggregates and achieve superior buffer subtraction. A co-flow system, in which the thin sample stream is sheathed with a matching buffer, enabled the use of high X-ray flux without causing protein radiation damage. This allowed us to obtain SAXS data even for variants that are marginally soluble. ^45, 46^ The aromatic variants showed small but consistent differences in their global dimensions and apparent scaling exponents as seen in a comparison of their raw data and normalized Kratky plots, respectively (Fig. S9, Fig. 4a). This is also true of variants where we alter the charge content and NCPR (Fig. S9, S10). We extracted apparent scaling exponents *ν*^app^ from the scattering data using an empirical molecular form factor ^47^. All apparent scaling exponents were below 0.5 (Table S3), implying that the variants adopt conformations with chain dimensions that are in the crossover regime between effective theta versus poor solvents.

**Fig. 4:**
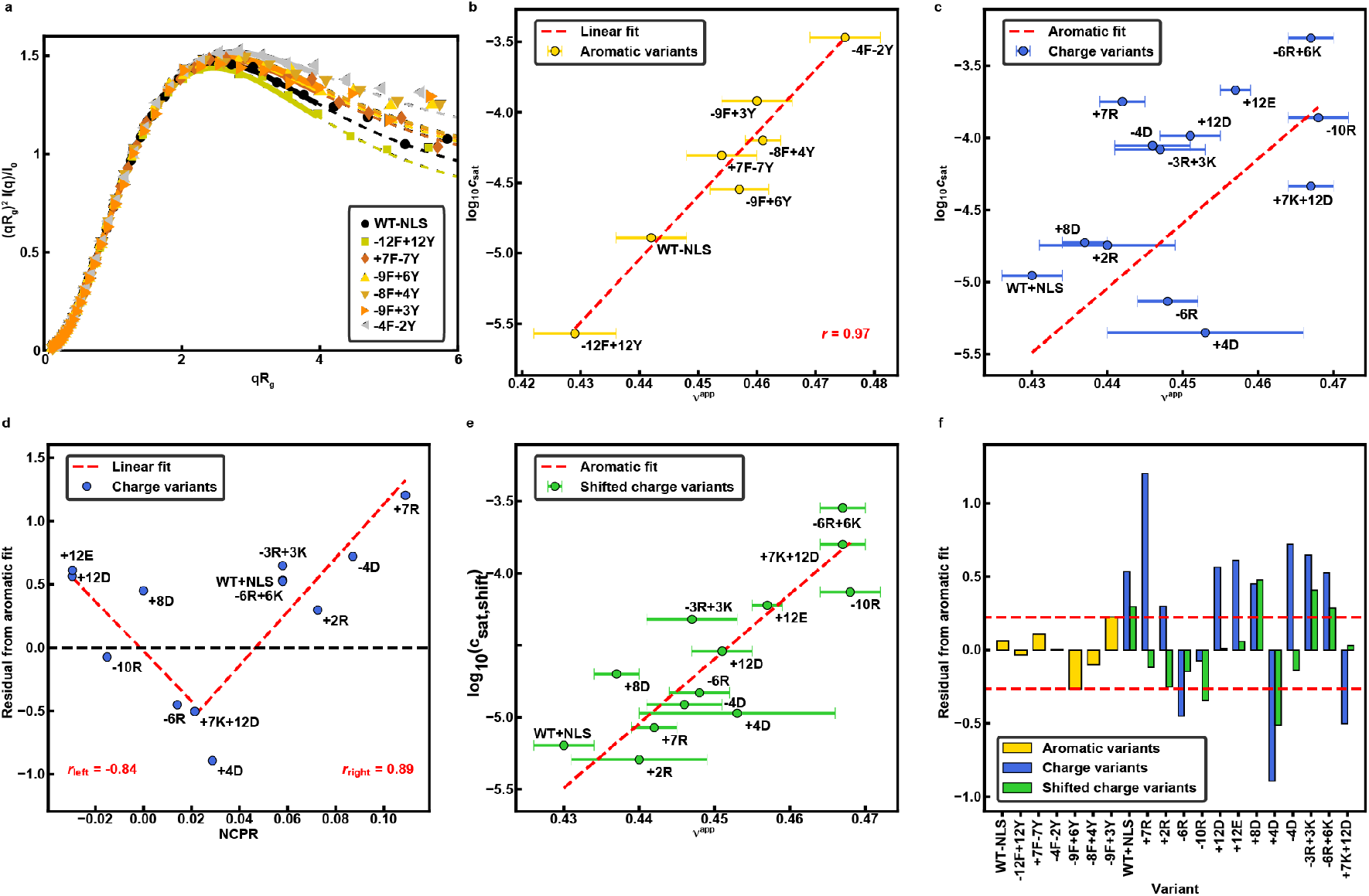
Mean-field electrostatic effects can disrupt the strong coupling between driving forces for single chain contraction and phase separation. **(a)** Normalized Kratky plot of SEC-SAXS data for aromatic variants of A1-LCD. Solid lines are fits to an empirical molecular form factor (MFF). Dashed lines show the predicted behavior at larger q values, where the experimental data are noisy. **(b)** Correlation between log_10_(*c*_sat_) and inferred values of *ν*^app^ for the aromatic variants. Red dashed line is the linear fit to the data. **(c)** Correlation between log_10_(*c*_sat_) and inferred *ν*^app^ values for the charge variants. The dashed line is the linear fit of the aromatic variants from (b). **(d)** Residuals from the correlation in (b) versus NCPR for the charge variants. Red dashed lines show two linear fits, i.e., for the variants whose NCPR is less than or equal to that of the +7K+12D variant and greater than or equal to that of the +7K+12D variant on the left and right, respectively. **(e)** Plot of log_10_(*c*_sat,shift_) against *ν*^app^ for the charge variants. Here, *c*_sat,shift_ refers to the *c*_sat_ values that are shifted based on the linear fits from (d). The dashed line is the linear fit of the aromatic variants from panel (b). **(f)** Residuals from the correlation in (b) for all variants. Data for charge variants are shown using unmodified and shifted *c*_sat_ values as in (c) and (e), respectively.

We analyzed the correlation between log_10_(*c*_sat_) and *ν*^app^ for the aromatic variants (Fig. 4b). These values show a strong positive correlation (Pearson *r*-value of 0.97) implying strong coupling behavior for aromatic variants even if the balance of Tyr and Phe residues varies. However, this correlation is abolished for the variants that alter net charge and the overall charge content (Fig. 4c). Instead, the data show that *c*_sat_ can be lowered or elevated by up to two orders of magnitude without altering ν^app^. Inspired by the results in Fig. 3b, which show that mean-field electrostatic effects can alter the driving force for phase separation, we undertook the following analysis: First, we used linear regression to extract parameters from Fig. 4b that quantify the correlation between log_10_(*c*_sat_) at 4°C and *ν*^app^ for the aromatic variants. We used these parameters to assess the correlation between log_10_(*c*_sat_) and *ν*^app^ for the different charge variants. Residuals from this linear regression analysis are shown in Fig. 4d. This analysis reveals a V-shaped profile when the residuals are plotted against NCPR. The implication is that mean-field electrostatic interactions can enable a decoupling of the driving forces for single-chain contraction and the driving forces for phase separation. Specifically, increases to NCPR can impact the collective interactions among many chains without having a significant impact on single chain dimensions as quantified by *ν*^app^. We accounted for the effects of mean-field electrostatic repulsions by rescaling the measured *c*_sat_ values according to the NCPR of each variant (see Methods) and asked if they fall on the line of best fit introduced in Fig. 4b. Indeed, NCPR-based rescaling of *c*_sat_ values lead to a recovery of the one-to-one correspondence between *ν*^app^ and *c*_sat_ (Fig. 4e,f). These results imply that the strong coupling behavior between single-chain and phase behavior can be disrupted by mean-field electrostatic effects. These interactions have to be weak enough to not influence single chain dimensions, and yet collectively strong enough to impact multichain phase behavior.

### Computational stickers-and-spacers model for simulations of phase separation

To formalize the connections between sequence features within PLCDs and overall phase behavior, we used LASSI, which is a lattice-based simulation engine for coarse-grained simulations of sequence- and / or architecture-specific phase transitions ^18^. In the current implementation, we use a single bead per residue version of the LASSI model. All interactions are isotropic, Tyr, Phe, and Arg are modeled as distinct stickers, and all other residues are weakly attracting spacers. Even Lys is modeled as a generic spacer, and therefore our adaptation of LASSI does not account for the three-body effects due to Lys-mediated interactions. Accordingly, we do not present results for the Lys variants.

Lattice energies for Tyr-Tyr, Tyr-Phe, Phe-Phe, Arg-Tyr/Phe, sticker-spacer, and spacer-spacer interactions were parameterized using a Gaussian process Bayesian optimization (GPBO) approach ^35^. GPBO was applied iteratively to explore parameter space and obtain energies by requiring that the single chain dimensions from coarse-grained simulations reproduce the apparent scaling exponents *ν*^app^ that we inferred from SAXS experiments (Fig. 5a). In dimensionless units, the optimized pairwise contact energies, inferred purely from comparisons of simulated and measured dimensions of single chains, range from -22 to -2 (Fig. 5b, Table S2). We use Metropolis Monte Carlo simulations to sample conformational / configurational space on the lattice ^18^. Accordingly, the transition probabilities for converting between pairs of conformations or configurations are proportional to exp(-ΔE/kT). Here, ΔE is the difference in energy between a pair of conformations / configurations. In the simulations, we set k = 1 and T is in the interval 30 ≤ T ≤ 60. Accordingly, in units of the dimensionless simulation temperature replacing Tyr-Tyr interactions with a Tyr-spacer interaction, which represents the largest change in ΔE, will range from ca. 0.33kT to 0.66kT, depending on the simulation temperature.

**Fig. 5:**
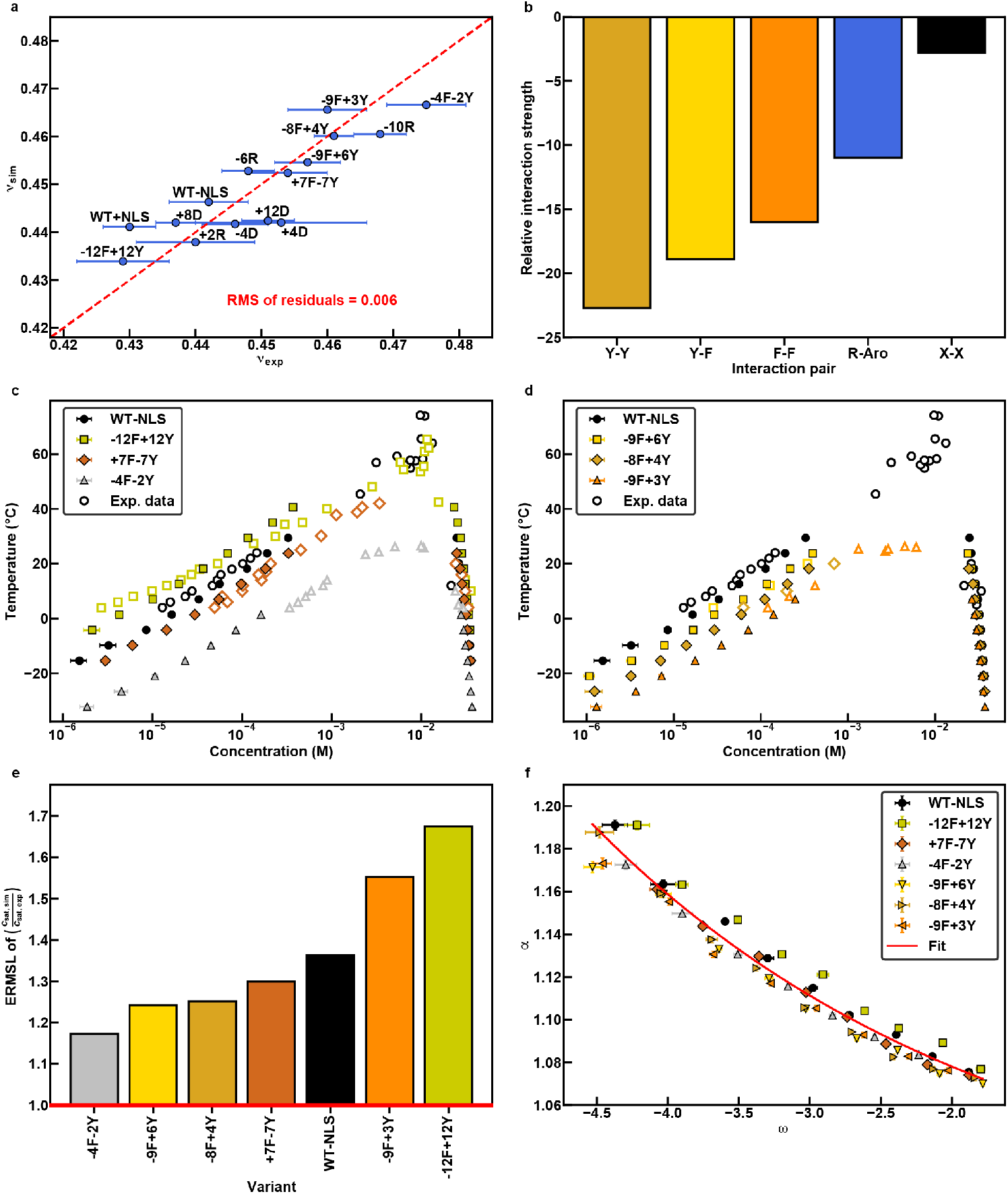
A computational stickers-and-spacers model captures the differences in sequence-binodal relationships for aromatic variants. **(a)** Correlation between *ν*^app^ from simulations, based on the optimized model, and inferences from measurements. Red dashed line indicates the target regime where the two sets of *ν*^app^ values are equal to one another. **(b)** Pairwise interaction strengths from the optimized computational model. **(c) – (d)** Computed (closed markers) and measured (open markers) binodals for aromatic variants. **(e)** Average factor of error of simulation-derived phase diagrams from experiment-derived phase diagrams of A1-LCD aromatic variants. A value of 1 indicates no error and a value of 10 indicates an error of an order of magnitude. **(f)** Plot of α against the width of the two-phase regime (ω) for aromatic variants. The red curve is a two-parameter exponential decay fit whereby α = 1 + exp[−*a* (ω−*b*)] and the parameters are *a* = 0.355 and *b* = −9.19.

The single-bead-per-residue LASSI model was used to model the phase behavior of all variants without changes to Lys residues, i.e., 18 variants. Binodals for aromatic variants were extracted from the simulation results and compared to those inferred from experimental measurements (Figs. 5c, 5d). Concentrations were converted from volume fractions to molar units and temperatures were converted from simulation units to degree Celsius using conversion factors introduced previously ^17^. We quantified the deviations between computed and measured binodals as the ratio of the computed and measured temperature-dependent *c*_sat_ values (Fig. 5e). The average error factor, computed across all variants, is 1.4, implying that on average the measured and calculated *c*_sat_ values are different by a factor of 1.4.

Building on accurate reproductions of the binodals for aromatic variants, we compared the differences in conformational preferences across phase boundaries. We start with the swelling ratio a ^48^, defined as the ratio of the average radius of gyration in the dense phase to that in the dilute phase. In Fig. 5f we plot a against the dimensionless width of the two-phase regime, defined as 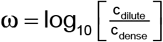 where *c*_dilute_ and *c*_dense_ are the dilute and dense phase concentrations at a specific temperature (Fig. S11). The swelling ratio approaches unity as *T* tends to the critical temperature (*T*_c_) (Fig. S11). We find that swelling ratios for all variants collapsed, without any adjustable parameters, onto a single master curve that could be fit to an exponential decay function (Fig. 5f). Accordingly, knowledge of the width of the two-phase regime allows us to infer the swelling ratio from the master curve. The implication is that if we supplement knowledge regarding the width of the two-phase regime with measurements of chain dimensions in the dilute phase, then the master curve allows us to infer dimensions of single chains in the dense phase. We find that PLCDs become more expanded in the dense phase when compared to the dilute phase. A swelling ratio of 1.2 translates to a dilution of intramolecular contacts by a factor of 1.8 within the dense phase and this points to the replacement of intramolecular interactions by intermolecular interactions.

Next, we applied our model to the subset of charge variants used in the parameterization (Fig. S12a,b) and to a set of charge variants that were not used in the parameterization (Fig. S12c). Note that the parameterization rests on the assumption of strong coupling between the driving forces for singlechain compaction and phase separation. Based on the results summarized in Fig. 4c-e it follows that the assumption of strong coupling gives rise to computed binodals that deviate significantly from measured binodals (Fig. S12a-c). Therefore, we built on insights from the analysis in Fig. 4c-e and included a meanfield NCPR-based adjustment to the potentials for simulations of multichain phase behavior. In these simulations, the pairwise interactions were weakened or strengthened by an amount that is proportional to the difference in NCPR values between that of the given variant and that of the WT (see SI Methods). Incorporation of this correction decreased the average error factor between the computed and measured binodals from 3.0 to 1.8 for the charge variants (Fig. 6a-c, Fig. S12d). These calculations provide support for the postulate that NCPR-mediated mean-field interactions can weaken the coupling between singlechain conformational equilibria and multichain phase equilibria.

**Fig. 6:**
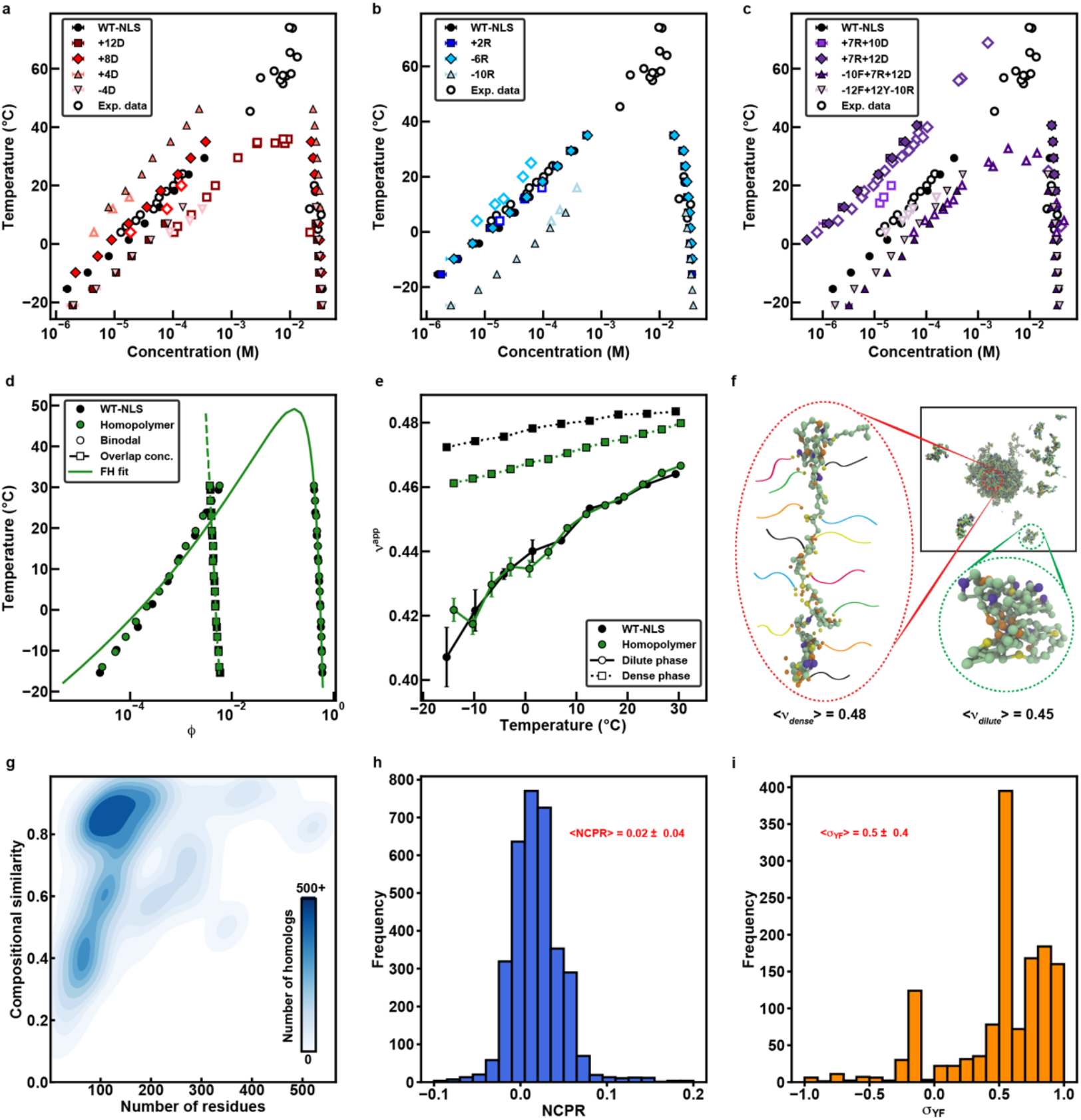
The stickers-and-spacers model can be generalized for charge variants and other IDPs. **(ac)** Computed (closed markers) and measured (open markers) binodals of charge variants. Simulations incorporate a mean-field NCPR-based term to capture the effects of charge variation. Variants in (c) were not used to parameterize the model and comparisons of computed and measured binodals provide a jack-knife testing of the accuracy of the computational model. **(d)** Computationally derived binodals (circular markers) and overlap concentrations (square markers) of WT A1-LCD and an equivalent homopolymer. The solid green line is a Flory-Huggins-based fit to the binodal for the equivalent homopolymer and the dashed green line is an extension of the overlap concentration line for this system. **(e)** Average *ν*^app^ values from dilute (circular markers; solid lines) and dense (square markers; dashed lines) phases of WT A1-LCD and its equivalent homopolymer. **(f)** Representative conformations for individual A1-LCD molecules drawn from the dense and dilute phases at 13°C. Beads are colored as follows: Tyr, yellow; Phe, orange; Arg, purple; all other residues, green. In the dense phase conformation, nearby stickers are shown with multi-colored tails to depict that the stickers originate from different protein chains. **(g)** 2D histogram quantifying the distributions of lengths for LCDs from FUS / FET family homologs and their compositional similarity to WT A1-LCD. **(h)** Distribution of NCPR values across LCDs from FUS / FET family homologs. **(i)** Distribution of Tyr versus Phe asymmetries for LCDs from FUS / FET family homologs.

### Distinguishing between stickers-and-spacers polymers and homopolymers

A stickers-and-spacers system is characterized by heterogeneity of interactions whereby sticker-sticker, sticker-spacer, and spacer-spacer interactions have distinct interaction strengths that depend on the types of stickers and spacers. Conversely, in a homopolymer model all interactions are equivalent. What can we learn from the mapping of stickers-and-spacers onto effective homopolymers? To answer this question, we parameterized an effective homopolymer model whose binodal overlays with that of WT A1-LCD (Fig. 6d). The resulting pairwise bead-to-bead interaction strengths in the equivalent homopolymer was -3.3. These pairwise energies are 1.2 times larger than spacer-spacer interactions in the stickers-and-spacers model and roughly seven times smaller than sticker-sticker interactions.

In polymer solutions, there exists a special concentration that equals the concentration of chain units within the pervaded volume of a single chain ^49^. This is known as the *overlap ConCentration c*\* - so named due to the high likelihood that chains overlap with one another when the solution concentration exceeds *c*\*. In dilute solutions (*c*<*c*\*), the properties of polymer solutions are governed exclusively by the interplay of intramolecular and chain-solvent interactions. Conversely, in semi-dilute solutions (*c*≈*c*\*), the physical properties of polymer solutions are governed by the multiway interplay among conformational fluctuations and their impact on intramolecular, intermolecular, and chain-solvent interactions. We used the mean end-to-end distance values in the single-chain limit ^38^ to compute temperature-dependent overlap volume fractions ϕ*(*T*) for the WT A1-LCD and its equivalent homopolymer. For temperatures below 20°C, ϕ_sat_(T) <ϕ*(T) i.e., the left arm of the binodal for A1-LCD is located to the left of the overlap line (Fig. 6d). Accordingly, for T < 20°C, the dispersed phase that coexists with the dense phase is a true dilute phase. However, we observe a crossover above ~20°C whereby ϕ_sat_(T) > ϕ*(T). Therefore, the dispersed phase that coexists with the condensate is actually semi-dilute for temperatures above 20 - 22°C.

The changes in conformational properties of PLCDs with sticker-and-spacer architectures across phase boundaries will be different from that of equivalent homopolymers. Cohesive, intramolecular inter-sticker interactions impose conformational strain on PLCD-like systems because the formation of sticker-sticker crosslinks comes at the expense of conformational restraints on weakly attracting spacer units that make up over 80% of the sequence. The strain that is built up in coexisting dilute / semi-dilute phases should be considerably different for the stickers-and-spacers model when compared to the effective homopolymer. Consequently, while the dilute phase single-chain dimensions are identical for the two models, the equivalent homopolymer is consistently more compact in the dense phase than the stickersand-spacers heteropolymer (Fig. 6e, Fig. S13).

Alleviation of strain should drive increased access to stickers for networks of intermolecular, inter-sticker crosslinks, and this is illustrated in Fig. 6f. A chain comprising *n_s_* stickers can engage in physical crosslinks with up to *n_s_* distinct chain molecules. This high networkability appears to be achievable in the dense phase as illustrated by excising a representative conformation from the dense phase and annotating it by the number of distinct chains that form inter-sticker crosslinks with the WT A1-LCD in its dense phase (Fig. 6f). These results highlight the importance of the stickers-and-spacers architecture and the heterogeneity of sticker versus spacer interactions for generating highly networked condensates - a defining feature of percolated network fluids ^19, 50^ whose material properties ^51^ will be governed by the timescales for making and breaking of physical crosslinks ^29^. In contrast, for equivalent homopolymers, the interactions are weak and uniform across the sequence. This leads to conformations with lower strains and a relative preference for intramolecular interactions that weaken the drive for chain expansion and intermolecular networking in dense phases.

## Discussion

In this work, we deployed a combination of methods to obtain systematic assessments of the driving forces for phase separation of PLCDs. Our results provide a molecular-level explanation for a range of prior observations while also uncovering new findings. These include the previously unappreciated context-dependent nature of Arg as an auxiliary sticker and the destabilizing effect of Lys due to three-body effects. In accord with predictions from theory and computations ^28^, we find that charged residues such as Asp and Glu destabilize phase separation due to their large effective solvation volumes. Our results are also congruent with extant data for other systems ^16, 52^, with Tyr being a stronger sticker than Phe. This congruence with findings for other systems suggests that our results are generalizable and transferable across a range of systems - a prospect made possible by the development of the mean-field model (Fig. 3b) and the ability to model sequence-specific phase behavior using LASSI (Figs. 5c, 5d, 6a-c).

Phase separation of A1-LCD variants is destabilized as NCPR increases either in the positive or negative direction. These results are congruent with recent reports for a very different low complexity domain from DDX4 ^33^. Interestingly, the optimum NCPR that maximizes the driving forces for phase separation of A1-LCD and designed variants thereof is ≈+0.02. We propose that the preference for excess positive charge might be due to a network of favorable cation-pi interactions, whereby the cations enable a delocalization of and favorable interactions with the pi clouds and ring systems ^53^. It could also be a reflection of the preferential condensation of solution anions around cationic residues ^54^. Testing the latter will require systematic assessments of the effects of the charge density, valence, and hydrophobicity of solution ions ^54, 55, 56^. Additionally, using a detailed assessment by SAXS measurements, we find that increases in NCPR weaken the coupling between the driving forces for single-chain compaction and phase separation. These results point to a route for tuning phase behavior via mean-field electrostatic effects, and is reminiscent of postulates pertaining to the contributions of mean-field polyelectrostatic effects to the regulation of fuzzy complexes by multisite phosphorylation of disordered proteins ^57^.

Are the rules extracted from our studies of A1-LCD variants transferable to PLCDs from other FUS / FET family proteins? To answer this question, we culled a large set of ~3,000 PLCDs from homologs of FUS / FET family proteins ^58, 59^. The pairwise compositional similarities between these sequences and A1-LCD were typically greater than 0.8. This suggests that the compositional biases uncovered for PLCDs across homologs of hnRNPA1 are preserved across PLCDs from homologs of FUS family proteins (Fig. 6g, Fig. S14). Interestingly, the mean NCPR was also found to be ≈+0.02 with a narrow dispersion about this value (Fig. 6h, Fig. S14). Overall, the preservation of compositional similarities suggests that the physicochemical rules that we have extracted here, combined with the computational model we have parameterized from our investigations of A1-LCD variants should be transferrable to large numbers of related PLCDs and enable proteome-level modeling of PLCD phase behavior.

Interestingly, unlike the PLCDs from homologs of hnRNPA1, the PLCDs from homologs of FUS / FET family proteins show a stronger bias toward Tyr over Phe (Fig. 6i). This suggests that the driving forces for condensate formation might be evolutionarily tuned by titrating σ_YF_ in PLCDs from homologs of hnRNPA1, whereas the tuning in PLCDs from homologs of FUS / FET family proteins is achieved by titrating the Tyr contents (Fig. S14). Accordingly, in condensates that encompass a diversity of PLCDs, the PLCD-specific variations in σ_YF_ and Tyr contents might enable the tuning of relative strengths of inter-sticker interactions across networks of heterotypic protein-protein interactions. Studies directed toward quantifying the implications of altering the interplay between networks of homotypic and heterotypic interactions, as controlled by variations in sequence-specific σ_YF_ values and fractions of Tyr residues, will be helpful for uncovering the physicochemical determinants of compositional control of condensates such as stress granules.

We also assessed the possibility that the broad distribution of σ_YF_ values for A1-LCD homologs is a reflection of covariations between Tyr / Phe contents and Gly / Ser contents. Indeed, analysis of PLCDs derived from 848 homologs of hnRNPA1 shows an interesting pattern of covariation (Fig. S15). Specifically, we observe positive correlations between the fractions of Gly and Tyr as well as the fractions of Ser and Phe, respectively. These in turn lead to a negative correlation between the fractions of Gly and Ser. The covariation trends suggest that the Gly / Ser content within PLCDs might vary in concert with the Tyr / Phe contents, providing a route to regulate the driving forces for phase separation and control material properties of condensates. The observed covariation trends point to hidden complexities that can be uncovered through large-scale assessments of the impact of covariations on PLCD phase behavior.

Surprisingly, the results in Fig. 6 suggest that the dispersed phases that coexist with condensates are likely to be in the semi-dilute regime for temperatures that are above ~ 20°C. This is important because unlike semi-dilute solutions, dilute solutions feature large-scale local concentration fluctuations of chain units that range from being near zero between polymers to considerably higher values due to conformational fluctuations of individual polymers ^49^. These local concentration fluctuations are considerably smaller in semi-dilute solutions ^49^. Accordingly, the semi-dilute nature of coexisting dispersed phases will enable an intrinsic buffering against concentration fluctuations at temperatures above ~20°C. Indeed, it is quite likely that dispersed phases that coexist with condensates in cells are in the semi-dilute regime due to macromolecular crowding and confinement effects. If true, this would have important implications for interfaces between dispersed and dense phases because in the semi-dilute / critical regime, the interfaces between coexisting phases are unlikely to be sharp and smooth. Instead, the interfaces will be defined by undulations that are on par with or larger than the dimensions of individual polymers ^60^. This will significantly lower the mean interfacial tension, a prediction that is in accord with ultra-low values estimated from microrheology experiments of model condensates in cells ^61, 62^. Importantly, in the semi-dilute / critical regime, the main determinants of selective permeabilities and transport properties of condensates will be governed by undulations of interfaces *viz.*, the wavelengths of capillary fluctuations ^60^ as opposed to interfacial tensions. Inasmuch as our *in vitro* observations have relevance to observations in cells, the connections we make here between criticality and the capillary properties of interfaces might provide an explanation for recent reports of seemingly unrestricted motions of protein factors across viral replication compartments in live cells ^63^. The general implications of our conjectures regarding the fluctuations of interfaces and their impact on selective permeability as well as transport properties will require further scrutiny.

## Methods

Details regarding the amino acid sequences of each designed variant, protein expression and purification, sample preparations for and collection of NMR, SAXS, and microscopy data are described in the Supplementary Information. Details of the bioinformatics analyses, design of LASSI simulations and the analysis of simulation results are also provided in the supplementary information.

### Measurements to construct binodals

Phase separation was induced by adding NaCl to a final concentration of 150 mM. Dilute and dense phase concentrations were determined as a function of temperature using a centrifugation method ^17, 39^. Briefly, phase separation of the sample was induced, and the dilute and dense phase were separated via centrifugation. Protein concentrations were determined by suitable dilutions of the samples and measurements of their absorbance at 280 nm. Points on the binodal above 30°C were determined via cloud point measurements using static light scattering as previously described^17, 40^. Briefly, phase separation of the sample was induced, and the dilute and dense phase were separated via centrifugation. Most of the dilute phase was removed and the remaining sample was heated above the critical point to enter the one-phase regime. 2 μL were removed with a positive displacement pipette and diluted to determine the protein concentration. 18 μL of the sample was transferred to a temperature equilibrated quartz DLS cuvette and the static light scattering was monitored as a function of temperature (temperature ramp 0.4°C/min) using a Wyatt NanoStar DLS instrument. The temperature at which the light scattering signal increases marks the transition of the one-phase regime into the two-phase regime. Cloud points at relatively lower temperatures/concentrations showed good agreement with coexistence points determined by centrifugation.

### van’t Hoff analysis

The chemical potentials of the dilute and dense phases can be written as 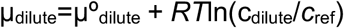 and 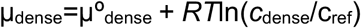, respectively. Here, 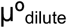 and 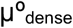 are the standard state chemical potentials of the PLCD molecules in the dilute and dense phases, respectively. Our measurements reveal that the concentrations of PLCD molecules in the dense phase show a weak dependence on temperature and on the sequence / compositions of the different variants. Accordingly, we set the reference concentration to be 0.03 M, which is also the approximate value of *c*_dense_ irrespective of temperature or the variant. This assumption is valid away from the critical point.

At equilibrium, μ_dilute_ = μ_dense_. Rearranging the equations above using the assumptions summarized thus far, we obtain the expression Δμ° = *RT*ln(*c*_dilute_/*c*_ref_), where 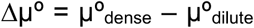. The difference in standard state chemical potentials quantifies the change in standard state free energies associated with transferring PLCD molecules from the dilute to dense phase. Assuming temperature independent values for standard state enthalpies (Δ*h*°) and entropies (Δ*s*°), we obtain the van’t Hoff equation ^64^ according to which:

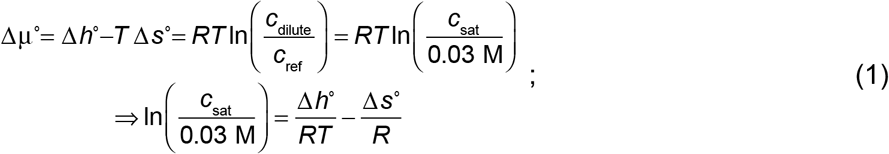

According to Equation (1), the slope of a plot of ln(*c*_sat_/*c*_ref_) versus (*RT*)^−1^, providing it is linear, will yield an estimate of the change in standard state enthalpy (Δ*h*°) and the intercept quantifies the standard state entropy (Δ*s*°) associated with transferring PLCD molecules across the phase boundary between coexisting dilute and dense phases. Fig. S8 shows the linear fits and associated parameters for all variants except +7R and +7R-12F+12Y. These variants were omitted because the binodals accessible via *in vitro* measurements are in the vicinity of the variant-specific critical points.

### Assessing the correlation between single-chain dimensions and c_sat_

In Figs. 4e, 4f, we shift the experimental *c*_sat_ values of the charge variants based on their associated NCPR values. This accounts for the greater sensitivity of *c*_sat_ on net charge. Shifted values of *c*_sat_ were obtained using parameters from the pair of linear fits in Fig. 4d. We then shift *c*_sat_ using log_10_(*c*_sat,shift_) =log_10_(*c*_sat_)-(*aq*+*b*). Here, *q* is the NCPR of the variant of interest. If the NCPR is less than 0.0214, which is the NCPR of the +7K+12D variant, the parameters are *a*= −19.7 and *b*= −0.0273. Alternatively, *a*= 21.1 and *b*= −0.989.

### LASSI simulations

Simulations were performed using LASSI, a lattice-based Monte Carlo engine for simulations of polymers with sticker-and-spacer architectures ^18^. Monte Carlo moves are accepted or rejected based on the Metropolis-Hastings criterion such that the probability of accepting a move is equal to min(1, exp(-βΔE)), where β= 1/kT, *T* is the simulation temperature, *k*= 1, and ΔE is the change in total system energy that accompanies the attempted move. Total system energies were calculated using a nearest neighbor model whereby any two beads that are within one lattice unit of each other along all three coordinate axes contribute to the total energy of the system. In our case, we define all pairwise interaction energies as absolute energies (Fig. 5b) that are scaled by the simulation temperature during the Metropolis-Hastings step. For example, the strongest interaction energy used in our simulations is -22.7 energy units and the range of simulation temperatures is 40-57 energy units. Within this temperature range, the strongest pairwise interaction is between -0.57kT and -0.40kT. Further details of the parameterization of the LASSI model, and the design and execution of single-chain and multi-chain simulations are provided in the Supplementary Information.

## Acknowledgments

We thank Furqan Dar for assistance with the LASSI simulations and Youlin Xia for help with NMR experiments. We are grateful to Furqan Dar, Alex Holehouse, Matthew King, Kiersten Ruff, J. Paul Taylor, and Xiangze Zeng for helpful discussions and critical reading of the manuscript. Microscopy images were acquired at the Cell & Tissue Imaging Center at SJCRH, which is supported by SJCRH and NCI (grant P30 CA021765). This work was supported by the US National Institutes of Health (grant 5R01NS056114 to RVP), the Air Force Office of Scientific Research (grant FA9550-20-1-0241 to R.V.P.), the St. Jude Collaborative Research Consortium on Membraneless Organelles (T.M. and R.V.P.), and the American Lebanese Syrian Associated Charities (to T.M.). Use of the Advanced Photon Source was supported by the U.S. Department of Energy under contract DE-AC02-06CH11357. LASSI is available for free download and use from https://github.com/Pappulab/LASSI and is released under the GPL - 3.0 license.

## Author contributions

A.B., M.F., R.V.P., and T.M. designed the study. A.B., W.M.B., I.P., and E.W.M. acquired different components of the experimental data and / or provided key reagents for experiments. M.F. and R.V.P. designed the computational and theoretical analysis and M.F. adapted and deployed the LASSI model. A.B., M.F., R.V.P., and T.M. wrote and revised multiple versions of the manuscript. All authors read and contributed revisions. R.V.P. and T.M. acquired funding.

## Additional information

Supplementary information is available with the online version of the paper. Correspondence and request for materials should be addressed to R.V.P. and T.M.

## References

1. Brangwynne CP, Eckmann CR, Courson DS, Rybarska A, Hoege C, Gharakhani J, et al. Germline P granules are liquid droplets that localize by controlled dissolution/condensation. Science 2009, 324(5935):1729–1732.

2. Li P, Banjade S, Cheng H-C, Kim S, Chen B, Guo L, et al. Phase transitions in the assembly of multivalent signalling proteins. Nature 2012, 483(7389):336–340.

3. Sabari BR, Dall’Agnese A, Boija A, Klein IA, Coffey EL, Shrinivas K, et al. Coactivator condensation at super-enhancers links phase separation and gene control. Science 2018, 361(6400):eaar3958.

4. Riggs CL, Kedersha N, Ivanov P, Anderson P. Mammalian stress granules and P bodies at a glance. Journal of Cell Science 2020, 133(16):jcs242487.

5. Yang P, Mathieu C, Kolaitis RM, Zhang P, Messing J, Yurtsever U, et al. G3BP1 Is a Tunable Switch that Triggers Phase Separation to Assemble Stress Granules. Cell 2020, 181(2):325–345 e328.

6. Sanders DW, Kedersha N, Lee DSW, Strom AR, Drake V, Riback JA, et al. Competing Protein-RNA Interaction Networks Control Multiphase Intracellular Organization. Cell 2020, 181(2):306–324.e328.

7. Guillen-Boixet J, Kopach A, Holehouse AS, Wittmann S, Jahnel M, Schlussler R, et al. RNA-Induced Conformational Switching and Clustering of G3BP Drive Stress Granule Assembly by Condensation. Cell 2020, 181(2):346–361 e317.

8. Mitchell SF, Jain S, She M, Parker R. Global analysis of yeast mRNPs. Nature Structural & Molecular Biology 2013, 20(1):127–133.

9. Cascarina SM, Elder MR, Ross ED. Atypical structural tendencies among low-complexity domains in the Protein Data Bank proteome. PLoS computational biology 2020, 16(1):e1007487.

10. Kim HJ, Kim NC, Wang Y-D, Scarborough EA, Moore J, Diaz Z, et al. Mutations in prion-like domains in hnRNPA2B1 and hnRNPA1 cause multisystem proteinopathy and ALS. Nature 2013, 495(7442):467–473.

11. Mackenzie IR, Nicholson AM, Sarkar M, Messing J, Purice MD, Pottier C, et al. TIA1 Mutations in Amyotrophic Lateral Sclerosis and Frontotemporal Dementia Promote Phase Separation and Alter Stress Granule Dynamics. Neuron 2017, 95(4):808–816 e809.

12. Schwartz JC, Cech TR, Parker RR. Biochemical Properties and Biological Functions of FET Proteins. Annual Review of Biochemistry 2015, 84(1):355–379.

13. Kwon I, Kato M, Xiang S, Wu L, Theodoropoulos P, Mirzaei H, et al. Phosphorylation-Regulated Binding of RNA Polymerase II to Fibrous Polymers of Low-Complexity Domains. Cell 2013, 155(5):1049–1060.

14. Molliex A, Temirov J, Lee J, Coughlin M, Kanagaraj AP, Kim HJ, et al. Phase separation by low complexity domains promotes stress granule assembly and drives pathological fibrillization. Cell 2015, 163(1):123–133.

15. Patel A, Lee Hyun O, Jawerth L, Maharana S, Jahnel M, Hein Marco Y, et al. A Liquid-to-Solid Phase Transition of the ALS Protein FUS Accelerated by Disease Mutation. Cell 2015, 162(5):1066–1077.

16. Wang J, Choi JM, Holehouse AS, Lee HO, Zhang X, Jahnel M, et al. A Molecular Grammar Governing the Driving Forces for Phase Separation of Prion-like RNA Binding Proteins. Cell 2018, 174(3):688–699 e616.

17. Martin EW, Holehouse AS, Peran I, Farag M, Incicco JJ, Bremer A, et al. Valence and patterning of aromatic residues determine the phase behavior of prion-like domains. Science 2020, 367(6478):694–699.

18. Choi J-M, Dar F, Pappu RV. LASSI: A lattice model for simulating phase transitions of multivalent proteins. PLoS computational biology 2019, 15(10):

19. Choi J-M, Holehouse AS, Pappu RV. Physical Principles Underlying the Complex Biology of Intracellular Phase Transitions. Annual Review of Biophysics 2020, 49:107–133.

20. Chong PA, Vernon RM, Forman-Kay JD. RGG/RG Motif Regions in RNA Binding and Phase Separation. Journal of Molecular Biology 2018, 430(23):4650–4665.

21. Vernon RM, Chong PA, Tsang B, Kim TH, Bah A, Farber P, et al. Pi-Pi contacts are an overlooked protein feature relevant to phase separation. eLife 2018, 7:e31486.

22. Brady JP, Farber PJ, Sekhar A, Lin Y-H, Huang R, Bah A, et al. Structural and hydrodynamic properties of an intrinsically disordered region of a germ cell-specific protein on phase separation. Proceedings of the National Academy of Sciences 2017, 114(39):E8194.

23. Nott Timothy J, Petsalaki E, Farber P, Jervis D, Fussner E, Plochowietz A, et al. Phase Transition of a Disordered Nuage Protein Generates Environmentally Responsive Membraneless Organelles. Molecular Cell 2015, 57(5):936–947.

24. Brangwynne CP, Tompa P, Pappu RV. Polymer physics of intracellular phase transitions. Nature Physics 2015, 11(11):899–904.

25. Alshareedah I, Kaur T, Ngo J, Seppala H, Kounatse L-AD, Wang W, et al. Interplay between Short-Range Attraction and Long-Range Repulsion Controls Reentrant Liquid Condensation of Ribonucleoprotein-RNA Complexes. Journal of the American Chemical Society 2019, 141(37):14593–14602.

26. Choi J-M, Hyman AA, Pappu RV. Generalized models for bond percolation transitions of associative polymers. Physical Review E 2020, 102:042403.

27. Harmon TS, Holehouse AS, Pappu RV. Differential solvation of intrinsically disordered linkers drives the formation of spatially organized droplets in ternary systems of linear multivalent proteins. New Journal of Physics 2018, 20(4):045002.

28. Harmon TS, Holehouse AS, Rosen MK, Pappu RV. Intrinsically disordered linkers determine the interplay between phase separation and gelation in multivalent proteins. eLife 2017, 6:e30294.

29. Jawerth L, Fischer-Friedrich E, Saha S, Wang J, Franzmann T, Zhang X, et al. Protein condensates as aging Maxwell fluids. Science 2020, 370(6522):1317.

30. Kumar S, Stecher G, Suleski M, Hedges SB. TimeTree: A Resource for Timelines, Timetrees, and Divergence Times. Molecular Biology and Evolution 2017, 34(7):1812–1819.

31. Greig JA, Nguyen TA, Lee M, Holehouse AS, Posey AE, Pappu RV, et al. Arginine-Enriched Mixed-Charge Domains Provide Cohesion for Nuclear Speckle Condensation. Molecular Cell 2020, 77(6):1237–1250.e1234.

32. Fisher RS, Elbaum-Garfinkle S. Tunable multiphase dynamics of arginine and lysine liquid condensates. Nature Communications 2020, 11(1):4628.

33. Crabtree MD, Holland J, Kompella P, Babl L, Turner N, Baldwin AJ, et al. Repulsive electrostatic interactions modulate dense and dilute phase properties of biomolecular condensates. bioRxiv 2020:2020.2010.2029.357863.

34. Frey S, Rees R, Schünemann J, Ng SC, Fünfgeld K, Huyton T, et al. Surface Properties Determining Passage Rates of Proteins through Nuclear Pores. Cell 2018, 174(1):202–217.e209.

35. Ruff KM, Harmon TS, Pappu RV. CAMELOT: A machine learning approach for coarse-grained simulations of aggregation of block-copolymeric protein sequences. The Journal of Chemical Physics 2015, 143(24):243123.

36. Dignon GL, Zheng W, Best RB, Kim YC, Mittal J. Relation between single-molecule properties and phase behavior of intrinsically disordered proteins. Proc Natl Acad Sci U S A 2018, 115(40):9929–9934.

37. Zeng X, Holehouse AS, Chilkoti A, Mittag T, Pappu RV. Connecting Coil-to-Globule Transitions to Full Phase Diagrams for Intrinsically Disordered Proteins. Biophysical Journal 2020, 119(2):402–418.

38. Wei MT, Elbaum-Garfinkle S, Holehouse AS, Chen CC, Feric M, Arnold CB, et al. Phase behaviour of disordered proteins underlying low density and high permeability of liquid organelles. Nature Chemistry 2017, 9(11):1118–1125.

39. Milkovic NM, Mittag T. Determination of Protein Phase Diagrams by Centrifugation. Methods Mol Biol 2020, 2141:685–702.

40. Peran I, Martin EW, Mittag T. Walking Along a Protein Phase Diagram to Determine Coexistence Points by Static Light Scattering. Methods Mol Biol 2020, 2141:715–730.

41. Martin EW, Holehouse AS. Intrinsically disordered protein regions and phase separation: sequence determinants of assembly or lack thereof. Emerging Topics in Life Sciences 2020, 4(3):307–329.

42. Dignon GL, Best RB, Mittal J. Biomolecular Phase Separation: From Molecular Driving Forces to Macroscopic Properties. Annu Rev Phys Chem 2020, 71:53–75.

43. Wang J, Choi J-M, Holehouse AS, Lee HO, Zhang X, Jahnel M, et al. A Molecular Grammar Governing the Driving Forces for Phase Separation of Prion-like RNA Binding Proteins. Cell 2018, 174(3):688–699.e616.

44. Crick SL, Ruff KM, Garai K, Frieden C, Pappu RV. Unmasking the roles of N- and C-terminal flanking sequences from exon 1 of huntingtin as modulators of polyglutamine aggregation. Proceedings of the National Academy of Sciences 2013, 110(50):20075.

45. Martin EW, Hopkins JB, Mittag T. Small angle x-ray scattering experiments of monodisperse samples close to the solubility limit. arXiv 2020:

46. Kirby N, Cowieson N, Hawley AM, Mudie ST, McGillivray DJ, Kusel M, et al. Improved radiation dose efficiency in solution SAXS using a sheath flow sample environment. Acta Crystallogr D Struct Biol 2016, 72(Pt 12):1254–1266.

47. Riback JA, Bowman MA, Zmyslowski AM, Knoverek CR, Jumper JM, Hinshaw JR, et al. Innovative scattering analysis shows that hydrophobic disordered proteins are expanded in water. Science 2017, 358(6360):238–241.

48. Muthukumar M. Thermodynamics of polymer solutions. The Journal of Chemical Physics 1986, 85(8):4722–4728.

49. Rubinstein M, Colby RH. Polymer Physics. Oxford University Press: New York, 2003.

50. Dias CS, Araújo NAM, Telo da Gama MM. Dynamics of network fluids. Advances in Colloid and Interface Science 2017, 247:258–263.

51. Roberts S, Harmon TS, Schaal JL, Miao V, Li K, Hunt A, et al. Injectable tissue integrating networks from recombinant polypeptides with tunable order. Nature Materials 2018, 17(12):1154–1163.

52. Dzuricky M, Rogers BA, Shahid A, Cremer PS, Chilkoti A. De novo engineering of intracellular condensates using artificial disordered proteins. Nature Chemistry 2020, 12(9):814–825.

53. Mahadevi AS, Sastry GN. Cation-π Interaction: Its Role and Relevance in Chemistry, Biology, and Material Science. Chemical Reviews 2013, 113(3):2100–2138.

54. Dahal YR, Schmit JD. Ion Specificity and Nonmonotonic Protein Solubility from Salt Entropy. Biophysical Journal 2018, 114(1):76–87.

55. Zheng W, Dignon GL, Jovic N, Xu X, Regy RM, Fawzi NL, et al. Molecular Details of Protein Condensates Probed by Microsecond Long Atomistic Simulations. The Journal of Physical Chemistry B 2020:

56. Boeynaems S, Bogaert E, Kovacs D, Konijnenberg A, Timmerman E, Volkov A, et al. Phase Separation of <em>C9orf72</em> Dipeptide Repeats Perturbs Stress Granule Dynamics. Molecular Cell 2017, 65(6):1044–1055.e1045.

57. Borg M, Mittag T, Pawson T, Tyers M, Forman-Kay JD, Chan HS. Polyelectrostatic interactions of disordered ligands suggest a physical basis for ultrasensitivity. Proceedings of the National Academy of Sciences 2007, 104(23):9650.

58. King OD, Gitler AD, Shorter J. The tip of the iceberg: RNA-binding proteins with prion-like domains in neurodegenerative disease. Brain Research 2012, 1462:61–80.

59. Dasmeh P, Wagner A. Natural Selection on the Phase-Separation Properties of FUS during 160 My of Mammalian Evolution. Molecular Biology and Evolution 2020:

60. Aarts DGAL, Schmidt M, Lekkerkerker HNW. Direct Visual Observation of Thermal Capillary Waves. Science 2004, 304(5672):847.

61. Brangwynne CP, Mitchison TJ, Hyman AA. Active liquid-like behavior of nucleoli determines their size and shape in Xenopus laevis oocytes. Proceedings of the National Academy of Sciences USA 2011, 108(11):4334–4339.

62. Feric M, Vaidya N, Harmon TS, Mitrea DM, Zhu L, Richardson TM, et al. Coexisting liquid phases underlie nucleolar subcompartments. Cell 2016, 165(7):1686–1697.

63. McSwiggen DT, Hansen AS, Teves SS, Marie-Nelly H, Hao Y, Heckert AB, et al. Evidence for DNA-mediated nuclear compartmentalization distinct from phase separation. eLife 2019, 8:e47098.

64. Holtzer A, Holtzer MF. Use of the van’t Hoff relation in determination of the enthalpy of micelle formation. The Journal of Physical Chemistry 1974, 78(14):1442–1443.

