## Supplemental Information for "Deciphering how naturally occurring sequence features impact the phase behaviors of disordered prion-like domains"

Includes details regarding the methods and analyses as well as Tables S1 – S3 and Supplemental Figures S1 – S15

### Supplemental Methods

*Details of constructs used in the current study*: All A1-LCD variants were based on the LCD (residues 186-320) from human hnRNPA1 (UniProt: P09651; Isoform A1-A). The coding sequences for the variants were synthesized (by Thermo Fisher or Genscript) including a coding sequence for an N-terminal ENLYFQGS TEV protease cleavage site and 5′ and 3′ attB sites for Gateway cloning. The sequences were recombined via LR reactions into the pDEST17 vector (Thermo Fisher), which includes an N-terminal 6xHis-tag coding sequence. In the expressed protein, the N-terminal 6xHis-tag was cleaved using the TEV protease cleavage site, leaving only an additional GS sequence at the N-terminus of each of the 28 constructs (underlined in Table S1). Amino acid sequence details of each of the constructs are shown in Table S1 below. In addition, variant Aro$^{-}$ from Martin et al. [1] was used and referred to as -4F-2Y.

**Table S1: Amino acid sequences of A1-LCD and designed variants**

| Construct | Amino acid sequence |
|---|---|
| A1-LCD$^{-NLS}$ | GSMASASSSQ RGRSGSGNFG GGRGGGFGGN DNFGRGGNFS GRGGFGGSRG GGGYGGSGDG YNGFGNDGSN FGGGGSYNDF GNYNNQSSNF GPMKGGNFGG RSSGGSGGGG QYFAKPRNQG GYGGSSSSSS YGSGRRF |
| A1-LCD$^{+NLS}$ | GSMASASSSQ RGRSGSGNFG GGRGGGFGGN DNFGRGGNFS GRGGFGGSRG GGGYGGSGDG YNGFGNDGSN FGGGGSYNDF GNYNNQSSNF GPMKGGNFGG RSSGPYGGGG QYFAKPRNQG GYGGSSSSSS YGSGRRF |
| A1-LCD$^{-4F-2Y}$ | GSMASASSSQ RGRSGSGNSG GGRGGGFGGN DNFGRGGNSS GRGGFGGSRG GGGYGGSGDG YNGFGNDGSN SGGGGSSNDF GNYNNQSSNF GPMKGGNFGG RSSGGSGGGG QYSAKPRNQG GYGGSSSSSS SGSGRRF |
| A1-LCD$^{-12F+12Y}$ | GSMASASSSQ RGRSGSGNYG GGRGGGYGGN DNYGRGGNYS GRGGYGGSRG GGGYGGSGDG YNGYGNDGSN YGGGGSYNDY GNYNNQSSNY GPMKGGNYGG RSSGGSGGGG QYYAKPRNQG GYGGSSSSSS YGSGRRY |
| A1-LCD$^{+7F-7Y}$ | GSMASASSSQ RGRSGSGNFG GGRGGGFGGN DNFGRGGNFS GRGGFGGSRG GGGFGGSGDG FNGFGNDGSN FGGGGSFNDF GNFNNQSSNF GPMKGGNFGG RSSGGSGGGG QFFAKPRNQG GFGGSSSSSS FGSGRRF |
| A1-LCD$^{-9F+6Y}$ | GSMASASSSQ RGRSGSGNFG GGRGGGYGGN DNYGRGGNYS GRGGFGGSRG GGGYGGSGDG YNGGGNDGSN YGGGGSYNDS GNYNNQSSNF GPMKGGNYGG RSSGGSGGGG QYGAKPRNQG |

| | |
|---|---|
| | GYGGSSSSS YGSGRRY |
| A1-LCD$^{-8F+4Y}$ | GSMASASSSQ RGRSGSGNFG GGRGGGYGGN DNGGRGGNYS GRGGFGGSRG GGGYGGSGDG<br>YNGGGNDGSN YGGGGSYNDS GNYNNQSSNF GPMKGGNYGG RSSGGSGGGG QYGAKPRNQG<br>GYGGSSSSS YGSGRRF |
| A1-LCD$^{-9F+3Y}$ | GSMASASSSQ RGRSGSGNFG GGRGGGYGGN DNGGRGGNYS GRGGFGGSRG GGGYGGSGDG<br>YNGGGNDGSN YGGGGSYNDS GNGNNQSSNF GPMKGGNYGG RSSGGSGGGG QYGAKPRNQG<br>GYGGSSSSS YGSGRRS |
| A1-LCD$^{-10R}$ | GSMASASSSQ GGSSGSGNFG GGGGGGFGGN DNFGGGGNFS GSGGFGGSGG GGGYGGSGDG<br>YNGFGNDGSN FGGGGSYNDF GNYNNQSSNF GPMKGGNFGG SSSGPYGGGG QYFAKPGNQG<br>GYGGSSSSS YGSGGGF |
| A1-LCD$^{-6R}$ | GSMASASSSQ GGRSGSGNFG GGRGGGFGGN DNFGGGGNFS GSGGFGGSRG GGGYGGSGDG<br>YNGFGNDGSN FGGGGSYNDF GNYNNQSSNF GPMKGGNFGG SSSGPYGGGG QYFAKPGNQG<br>GYGGSSSSS YGSGGRF |
| A1-LCD$^{+2R}$ | GSMASASSSQ RGRSGSGNFG GGRGGGFGGN DNFGRGGNFS GRGGFGGSRG GGGYGGSGDG<br>YNGFRNDGSN FGGGGRYNDF GNYNNQSSNF GPMKGGNFGG RSSGPYGGGG QYFAKPRNQG<br>GYGGSSSSS YGSGRRF |
| A1-LCD$^{+7R}$ | GSMASASSSQ RGRSGRGNFG GGRGGGFGGN DNFGRGGNFS GRGGFGGSRG GGRYGGSGDR<br>YNGFGNDGRN FGGGGSYNDF GNYNNQSSNF GPMKGGNFRG RSSGPYGRGG QYFAKPRNQG<br>GYGGSSSRS YGSGRRF |
| A1-LCD$^{-2K}$ | GSMASASSSQ RGRSGSGNFG GGRGGGFGGN DNFGRGGNFS GRGGFGGSRG GGGYGGSGDG<br>YNGFGNDGSN FGGGGSYNDF GNYNNQSSNF GPMGGGNFGG RSSGPYGGGG QYFAGPRNQG<br>GYGGSSSSS YGSGRRF |
| A1-LCD$^{-3R+3K}$ | GSMASASSSQ RGKSGSGNFG GGRGGGFGGN DNFGRGGNFS GRGGFGGSKG GGGYGGSGDG<br>YNGFGNDGSN FGGGGSYNDF GNYNNQSSNF GPMKGGNFGG RSSGGSGGGG QYFAKPRNQG<br>GYGGSSSSS YGSGRKF |
| A1-LCD$^{-6R+6K}$ | GSMASASSSQ KGKSGSGNFG GGRGGGFGGN DNFGKGGNFS GRGGFGGSKG GGGYGGSGDG<br>YNGFGNDGSN FGGGGSYNDF GNYNNQSSNF GPMKGGNFGG KSSGGSGGGG QYFAKPRNQG<br>GYGGSSSSS YGSGRKF |
| A1-LCD$^{-10R+10K}$ | GSMASASSSQ KGKSGSGNFG GGKGGGFGGN DNFGKGGNFS GKGGFGGSKG GGGYGGSGDG<br>YNGFGNDGSN FGGGGSYNDF GNYNNQSSNF GPMKGGNFGG KSSGGSGGGG QYFAKPKNQG<br>GYGGSSSSS YGSGKKF |
| A1-LCD$^{-4D}$ | GSMASASSSQ RGRSGSGNFG GGRGGGFGGN GNFGRGGNFS GRGGFGGSRG GGGYGGSGGG<br>YNGFGNSGSN FGGGGSYNGF GNYNNQSSNF GPMKGGNFGG RSSGPYGGGG QYFAKPRNQG<br>GYGGSSSSS YGSGRRF |
| A1-LCD$^{+4D}$ | GSMASASSSQ RDRSGSGNFG GGRGGGFGGN DNFGRGGNFS GRGDFGGSRG GGGYGGSGDG<br>YNGFGNDGSN FGGGGSYNDF GNYNNQSSNF GPMKGGNFGG RSSDPYGGGG QYFAKPRNQG<br>GYGGSSSSS YDSGRRF |
| A1-LCD$^{+8D}$ | GSMASASSSQ RDRSGSGNFG GGRDGGFGGN DNFGRGDNFS GRGDFGGSRD GGGYGGSGDG<br>YNGFGNDGSN FGGGGSYNDF GNYNNQSSNF GPMKGGNFGG RSSDPYGGGG QYFAKPRNQD<br>GYGGSSSSS YDSGRRF |
| A1-LCD$^{+12D}$ | GSMASADSSQ RDRDDSGNFG DGRGGGFGGN DNFGRGGNFS DRGGFGGSRG DGGYGGDGDG<br>YNGFGNDGSN FGGGGSYNDF GNYNNQSSNF DPMKGGNFGD RSSGPYDGGG QYFAKPRNQG<br>GYGGSSSSS YGSDRRF |
| A1-LCD$^{+12E}$ | GSMASAESSQ REREESGNFG EGRGGGFGGN DNFGRGGNFS ERGGFGGSRG EGGYGGEGDG<br>YNGFGNDGSN FGGGGSYNDF GNYNNQSSNF EPMKGGNFGE RSSGPYEGGG QYFAKPRNQG<br>GYGGSSSSS YGSERRF |
| A1-LCD$^{+7R+10D}$ | GSMASADSSQ RDRDGRGNFG DGRGGGFGGN DNFGRGGNFS DRGGFGGSRG GGRYGGDGDR<br>YNGFGNDGRN FGGGGSYNDF GNYNNQSSNF DPMKGGNFRD RSSGPYDRGG QYFAKPRNQG<br>GYGGSSSRS YGSDRRF |
| A1-LCD$^{+7R+12D}$ | GSMASADSSQ RDRDDRGNFG DGRGGGFGGN DNFGRGGNFS DRGGFGGSRG DGRYGGDGDR<br>YNGFGNDGRN FGGGGSYNDF GNYNNQSSNF DPMKGGNFRD RSSGPYDRGG QYFAKPRNQG<br>GYGGSSSRS YGSDRRF |
| A1-LCD$^{+7K+12D}$ | GSMASADSSQ RDRDDKGNFG DGRGGGFGGN DNFGRGGNFS DRGGFGGSRG DGKYGGDGDK<br>YNGFGNDGKN FGGGGSYNDF GNYNNQSSNF DPMKGGNFKD RSSGPYDKGG QYFAKPRNQG<br>GYGGSSSKS YGSDRRF |
| A1-LCD$^{-2R-2K+3D}$ | GSMASASSSQ DGRSGSGNFG GGRGGGFGGN DNFGRGGNFS GRGGFGGSRG GGGYGGSGDG<br>YNGFGNDGSN FGGGGSYNDF GNYNNQSSNF GPMDGGNFGG RSSGPYGGGG QYFADPRNQG<br>GYGGSSSSS YGSGGRF |
| A1-LCD$^{-4R-2K+5D}$ | GSMASASSSQ DGRSGSGNFG GGDGGGFGGN DNFGRGGNFS GGGGFGGSRG GGGYGGSGDG<br>YNGFGNDGSN FGGGGSYNDF GNYNNQSSNF GPMDGGNFGG RSSGPYGGGG QYFADPRNQG |

| | |
|---|---|
| | GYGGSSSSSS YGSGDRF |
| A1-LCD$^{+7K+12D}$ blocky | GSMASAKSSQ RDRDDDGNFG KGRGGGFGGN KNFGRGGNFS KRGGFGGSRG KGKYGGKGDD YNGFGNDGDN FGGGGSYNDF GNYNNQSSNF DPMDGGNFDD RSSGPYDDGG QYFADPRNQG GYGGSSSSKS YGSKRRF |
| A1-LCD$^{-12F+12Y-10R}$ | GSMASASSSQ GGSSGSGNYG GGGGGGYGGN DNYGGGGNYS GSGGYGGSGG GGGYGGSGDG YNGYGNDGSN YGGGGSYNDY GNYNNQSSNY GPMKGGNYGG SSSGPYGGGG QYYAKPGNQG GYGGSSSSSS YGSGGGY |
| A1-LCD$^{-10F+7R+12D}$ | GSMASADSSQ RDRDDRGNFG DGRGGGGGGN DNFGRGGNGS DRGGGGGSRG DGRYGGDGDR YNGGGNDGRN GGGGGSYNDG GNYNNQSSNG DPMKGGNGRD RSSGPYDRGG QYGAKPRNQG GYGGSSSSRS YGSDRRG |

*Protein expression and purification*: All hnRNPA1-LCD variants were expressed in *E. coli* BL21-Gold (DE3) strain in ZYM5052 auto induction media at 37°C for 24 hours. For NMR samples, cultures were grown in isotopically labeled M9 media, induced at $OD_{600}$=0.8 with 1 mM IPTG and cultured at 37°C for an additional 6 hours. Cell pellets were resuspended in 50 mM MES pH 6.0, 500 mM NaCl, 20 mM 2-mercaptoethanol and lysed via sonication. Cell lysates were centrifuged, and the variants were purified from insoluble inclusion bodies as previously described [1]. The inclusion bodies were resuspended in 6 M GdmHCl, 20 mM Tris pH 7.5, 15 mM imidazole overnight at 4°C. Solutions of solubilized inclusion bodies were cleared by centrifugation, and supernatants were loaded onto self-packed columns of chelating Sepharose fast flow beads (GE Healthcare) charged with nickel sulfate. The columns were washed with 4 column volumes of 4 M urea, 20 mM Tris pH 7.5, 15 mM imidazole. Proteins were eluted from the Ni-NTA resin with 4 M urea, 20 mM Tris pH 7.5, 500 mM imidazole. TEV cleavage of the 6xHis-tag was done in 2 M urea, 20 mM Tris pH 7.5, 50 mM NaCl, 0.5 mM EDTA, 1 mM DTT overnight at 4°C. Cleaved protein solutions were loaded onto Ni-NTA columns. The flow-through and wash fractions were collected and concentrated using a 3000 MWCO Amicon centrifugal filter. As a final purification step, the samples were passed in 2 M GdmHCl, 20 mM MES pH 5.5 over a S75 Superdex size exclusion column (GE Healthcare). The identity of each protein was confirmed via intact mass spectrometry. All proteins were stored in 4 M GdmHCl, 20 mM MES pH 5.5 at 4°C. For the -2R-2K+3D construct, the procedure was modified as follows. The sample was cleaved in a minimum of 30 mL of buffer per 1 L of culture. Following the post-cleavage nickel column, the sample was rapidly exchanged into 20 mM MES pH 5.5 and 6 M GdmHCl using a 10K MWCO 15 mL Amicon centrifugal filter prior to size exclusion by a Superdex 75 column to avoid the protein being concentrated at a pH near its theoretical pI. For the -4R-2K+5D construct the procedure was modified such that the protein was cleaved in a minimum of 30 mL of buffer per 1 L of culture. The sample pH was then rapidly increased after the post-cleave nickel column by adding 1/10 volume of 1 M CAPS pH 10.5 before concentrating with a 10K MWCO 15 mL Amicon centrifugal filter. The sample was then subjected to size exclusion chromatography using a Superdex 75 column equilibrated with 20 mM CAPS pH 10.5, 2 M GdmHCl.

*Buffer exchange to remove denaturant*: Buffer exchange was achieved in two-steps. First, the protein in 4 M GdmHCl, 20 mM MES pH 5.5 storage buffer was exchanged into 1 M MES pH 5.5 by multiple dilution and concentration steps using a 3K MWCO Amicon centrifugal filter as previously described [1]. The protein was then dialyzed overnight against 20 mM HEPES pH 7.0 (without excess salt) at room temperature. The pH of the buffer was adjusted using ammonium hydroxide to prevent the introduction of excess salt into the sample. The protein was filtered through a 0.22 mm Millex-GV filter (Merck) to remove potential aggregates from the solution, which might have formed during dialysis.

*Measurements of saturation concentrations for specific variants that required special handling*: Experiments on variant +8D was carried out in 20 mM HEPES, 150 mM NaCl pH 8.0 because of its net neutral charge. Because +7F-7Y lacks Tyr residues, its protein concentration was determined at 205 nm using a Cary 300 UV-Vis spectrophotometer (Agilent). For determination of low protein concentrations, a 10 mm pathlength quartz cuvette was used.

We also measured saturation concentrations for variant +7K+12D as a function of pH. The sample of variant +7K+12D in denaturing buffer was rapidly exchanged into non-denaturing buffers using Zeba spin columns (Thermo Fischer) following standard procedures. The columns were equilibrated with buffers prepared at room temperature to contain 150 mM NaCl and 20 mM of one of the following buffering agents, MES at pH 5.5, and 6.5, HEPES at pH 6.5, 7, and 8, Tris at pH 8, and 9, and HEPBS at pH 8, 8.3, 8.7, and 9. The pH of each buffering condition was measured at 4ºC to account for temperature dependent $pK_a$ shifts of the buffer. The saturation concentration at 4ºC was then measured by separating dilute and dense phase by centrifugation as described in the Methods. All measurements were done as at least 3 replicates. The theoretical protein net charge at each pH was calculated using protpi.ch. The protonated state of the lysine side chain at each pH was calculated using the estimated net charge of the protein and accounting for the charge state of the amino terminal at the given pH, assuming the remainder of the charge difference results from deprotonated Lys residues.

*Small Angle X-ray Scattering* (*SAXS*) *measurements*: All measurements were performed at BioCat (beamline 18ID at the Advanced Photon Source, Chicago) with in-line size exclusion chromatography (SEC-SAXS) as previously reported [1, 2]. Experiments were conducted at room temperature in 20 mM HEPES, 150 mM NaCl, pH 7.0. Protein samples stored in 4 M GdmHCl, 20 mM MES pH 5.5 were loaded onto either a Superdex 75 5/150 GL or a Superdex 75 Increase 10/300 column (GE Life Science) with a flow rate of 0.4 mL/min. The column eluent passed through the UV monitor and proceeded through the SAXS sheath flow capillary in the coflow system [3]. Scattering intensity was recorded using a Pilatus3 1 M (Dectris) detector placed 3.5 m from the sample providing a q-range of 0.004-0.4 $Å^{-1}$. Exposure time was 0.5 sec. Raw SAXS data was reduced at the beamline using BioXTAS RAW 1.6.3 and 2.0.2 [4]. Buffer subtraction, Guinier fits, and Kratky transformations were performed using the BioXTAS Raw software [4]. Raw data were additionally fit using an empirically derived molecular form factor (MFF) developed by Riback et al. [5].

*Microscopy*: Differential interference contrast microscopy (DIC) images were obtained at room temperature using a Nikon Eclipse Ni Widefield microscope with a 20X objective. Samples were prepared by adding NaCl to 150 mM to the protein stock solution. Protein concentrations were slightly above their corresponding $c_{sat}$ at 20°C (see Figure 2c-g). 2 μL of the protein solution was sandwiched between two coverslips sandwiched with 3M 300 LSE high-temperature double-sided tape (0.34 mm) with a window for microscopy cut out.

*NMR spectroscopy*:

NOESY and TOCSY experiments for A1-LCD Δhexa were acquired on either a Bruker Avance 1.1 GHz or 850 MHz spectrometer equipped with TCI triple-resonance cryogenic probes and pulse-field gradient units. A $^{13}C$-resolved $^{1}H^{aromatic}$-$^{1}H^{aliphatic}$ NOESY spectrum (64 scans, 2048 ($^{1}H$) × 64 ($^{13}C$) × 80 ($^{1}H$) complex data points, with 14.2 ppm, 16.0 ppm, and 1.5 ppm as $^{1}H$, $^{13}C$ and $^{1}H$ sweep width, respectively) and a mixing time of 250 ms were measured at 1.1 MHz and 313 K in 20 mM HEPES, 200 mM NaCl, 5 mM TCEP, 150 μM DSS, and 5% D2O at pH 6.3. $^{13}C$ editing was not necessary for aromatic protons as they are well separated, but all post-NOE protons were $^{13}C$ resolved. The concentration of the Δhexa LCD was ~800 μM. 2D planes from the 3D spectra corresponding to the arginine δ-position were assessed for NOEs between the arginine δ-proton and the aromatic protons. A triple resonance (H)CC(CO)NH spectrum (16 scans, 2048 ($^{1}H$) × 80 ($^{13}C$) × 200 ($^{1}H$) complex data points, with 16.3 ppm, 22.0 ppm, and 54.0 ppm as the $^{1}H$, $^{13}C$ and $^{1}H$ sweep width, respectively) was used to assign the arginine sidechain frequencies. Data were processed using BRUKER Topspin version 4.0, NMRPipe version 10.4 [6] and analyzed using NMRfam SPARKY [7]. All spectra were referenced directly using DSS for the $^{1}H$ dimension; $^{13}C$ and $^{15}N$ frequencies were referenced indirectly.

NMR data for A1-LCD +7K+12D were acquired on Bruker Avance 600 and 800 MHz spectrometers equipped with TCI triple-resonance cryogenic probes and pulsed-field gradient units. All samples were prepared in a buffer consisting of 20 mM HEPES pH 6.8, 0.5 mM EDTA and 10% $D_2O$ at 20°C. For assignment, samples of $^{15}N$,$^{13}C$ +7K+12D with concentrations between 85 and 220 μM were used to acquire standard triple-resonance backbone assignment experiments based on a sensitivity enhanced $^{1}H$-$^{15}N$ HSQC (32 scans, 2048 x 512 complex data points, with 12 ppm and 20 ppm as $^{1}H$ and $^{15}N$ sweep widths). These included HNCACB and CBCA(CO)NH (16 scans, 2048 ($^{1}H$) × 64 ($^{15}N$) × 128 ($^{13}C$) complex data points, with 12 ppm, 20 ppm, and 72 ppm as $^{1}H$, $^{15}N$ and $^{13}C$ sweep width, respectively), HN(CA)CO (16 scans, 2048 ($^1$H) × 64 ($^{15}$N) × 128 ($^{13}$C) complex data points, with 10 ppm, 20 ppm, and 72 ppm as $^1$H, $^{15}$N and $^{13}$C sweep widths, respectively), HNCO (8 scans, 2048 ($^1$H) × 64 ($^{15}$N) × 80 ($^{13}$C) complex data points, with 10 ppm, 20 ppm, and 14 ppm as $^1$H, $^{15}$N and $^{13}$C sweep widths, respectively), and NH(CA)NNH (32 scans, 2048 ($^1$H) × 50 ($^{15}$N ) × 100 ($^{15}$N) complex data points, with 10 ppm, 20 ppm, and 22 ppm as $^1$H, $^{15}$N F1 and $^{15}$N F2 sweep widths, respectively) spectra.

Data were processed using BRUKER Topspin version 3.2, or NMRPipe (v.7.9) [6] and analyzed using NMRviewJ [8]. All spectra were referenced directly using DSS for the $^1$H dimension, $^{13}$C and $^{15}$N frequencies were referenced indirectly.

$^{15}$N $R_2$ relaxation experiments were acquired at 600 MHz at 293 K using a standard Carr-Purcell-Meiboom-Gill (CPMG)-based Bruker pulse program (32 scans, 2048 ($^1$H) x 256 ($^{15}$N) complex data points) with the following delays of 16.8, 33.5, 67, 100.5, 134.1, 167.6, 251.4, and 335.2 ms with a recovery delay of 3 s. Due to significant overlap in the spectra Peakipy was used to attempt to deconvolute the peak intensities. The relaxation rates for residues 13, 15, 17, 19, 20, 21, 23, 25, 26, 27, 29, 31, 36, 37, 38, 39, 40, 42, 47, 51, 52, 53, 56, 57, 58, 59, 60, 61, 62, 66, 67, 68, 71, 74, 77, 78, 83, 86, 88, 89, 90, 93, 96, 97, 98, 100, 103, 110, 121, 122, 124, 125, 126, 127, 130, 131, and 132 were omitted as the overlap was too great for deconvolution. Fitting of the $R_2$ rate profile to an analytical model was done as previously described [1]. To account for gaps in experimental data, the maximum cluster height was limited to 9 $s^{-1}$.

*Bioinformatics analysis*: Homologous sequences were identified using version 5.0 of the EggNOG database [9]. Sequences of homologs were aligned using the EMBL-EBI Clustal Omega tool [10]. The sequences were then trimmed to include only intrinsically disordered regions using the UniProt annotation [11] of the canonical human isoform. Sequence analyses were performed using the localCIDER tool [12]. The sequences culled for LCDs from homologs of hnRNPA1 and FUS / FET family proteins are included in separate Supplementary spreadsheets.

*Analysis of sequence compositions*: Pairwise compositional similarities were determined in the following manner: (1) For each sequence, we create a 20×1 compositional vector. Each vector is of the form: ($f_A$, $f_C$, … $f_Y$). Here, each element quantifies the fraction of each amino acid within a sequence. (2) For a pair of compositional vectors we compute the dot product of the two vectors. (3) Next, we divide the dot product by the product of the magnitudes of the vectors. This gives a value between 0 and 1, where values closer to 1 indicate higher compositional similarity. In Fig. 1c-g, the sequences analyzed share a compositional similarity with the WT sequence of at least 0.8 (770 sequences).

*Analysis of $c_{sat}$ data using a mean-field, stickers-and-spacers model*: Wang et al. [13] adapted the mean-field stickers and spacers model of Semenov and Rubinstein [14] to a system with $n_A$ stickers of type A and $n_B$ stickers of type B. In this model, the saturation concentration $c_{sat}$ was shown to be proportional to $(n_A n_B)^{-1}$ providing the heterotypic interactions among A and B stickers are the only determinants of $c_{sat}$. The model of Wang et al. [13] was generalized by Choi et al. [15] to account for the competing effects of homotypic A-A and B-B interactions. Additionally, Choi et al., accounted for cooperative effects, whereby the strengths of inter-sticker interactions can either be enhanced or weakened due to the influence of three-body interactions on the strengths of inter-sticker interactions. We adapted the approach of Choi et al. [15] to obtain rescaled values of $c_{sat}$ that were analyzed as a function of NCPR. Here, we choose the aromatic residues (Tyr / Phe) as the primary stickers and Arg as auxiliary stickers.

In the generalized mean-field model of Choi et al. [15], $c_{sat}$ is governed by the numbers of aromatic residues ($n_a$), the numbers of Arg residues ($n_R$), and the strengths of inter-aromatic ($\lambda_{aa}$) and aromatic-Arg ($\lambda_{aR}$) interactions. Note that the λ-values are dimensionless quantities. Accordingly, the functional form for $c_{sat}$, written in terms of a multiplicative constant is as shown in Equation (1).

$$c_{sat} = k\left(\lambda_{aa} n_a^2 + 2\lambda_{aR} n_a n_R\right); \quad (1)$$

Here, $k$ is a constant that converts the right-hand side into units of concentrations. The value for $k$ can be extracted by linear regression of measured $c_{sat}$ values plotted against the quantity in the parenthesis on the right-hand side of Equation (1) [13]. In our analysis, we focus on a rescaling of $c_{sat}$ and therefore we do not need an estimate for $k$. The first step in the rescaling, which accounts for the contributions of aromatic and Arg residues as stickers is written as:

$$c'_{sat} = c_{sat}\left(\lambda_{aa} n_a^2 + 2\lambda_{aR} n_a n_R\right); \tag{2}$$

Note that we set $\lambda_{aa} = 1$ and hence the only free parameter in the regression analysis is the value of $\lambda_{aR}$. If the only determinants of $c_{sat}$ were inter-sticker interactions, then the expectation would be that, with appropriate parameterization of $\lambda_{aR}$, the values of $c'_{sat}$ would be similar to one another for all A1-LCD variants. Instead, we observe the emergence of a V-shaped profile for $c'_{sat}$ plotted against NCPR, especially for the Arg and Asp/Glu variants (Fig. S2c).

We also notice that the Lys variants have considerably higher $c'_{sat}$ values than would be expected based on the numbers of aromatic and Arg residues in these variants. The implication is that Lys plays a distinctive role, not just as a high-excluded volume spacer, but also in terms of its impact on the strengths of inter-sticker interactions. This model emerges from findings regarding the influence of positive and / or negative cooperativity on pi-pi and cation-pi interactions [16]. Here, we reason, based on the model of Choi et al. [15] that Lys residues appear to weaken inter-sticker interactions via three-body interactions. This effect is captured in Equation (3) as:

$$c''_{sat} = c_{sat}\left[n_a^2\left(\lambda_{aa} - 3\lambda_K n_K\right) + n_a n_R\left(2\lambda_{aR} - 6\lambda_K n_K\right)\right]; \tag{3}$$

Here, $\lambda_K$ quantifies the extent to which Lys residues impact the effective strengths of inter-sticker interactions and $n_K$ is the number of protonated Lys residues. Notice that the effects of Lys residues are incorporated as contributions that affect $c_{sat}$ via three-body interactions. The impact of accounting for the destabilizing effects of Lys residues is summarized in Fig. S2d. This requires parameterization of $\lambda_{aR}$ and $\lambda_K$. This two-parameter fit of Equation (3) shows that the rescaled $c_{sat}$ values at 4˚C collapse onto the V-shape profile that is plotted against NCPR (Fig. S2d).

Finally, since we have measurements of $c_{sat}$ at a series of different temperatures, we account for the temperature dependence by noting that the dilute arms of the binodals are linear on a semi-log scale implying that the temperature dependence of $c_{sat}$ may be written as:

$$c_{sat}(T) = c_{sat}(T_0)\exp\left[-\left(\frac{T - T_0}{m}\right)\right]; \tag{4}$$

Here, $c_{sat}(T_0)$ is the $c_{sat}$ value at the reference temperature of 277 K and $T$ is the actual temperature at which $c_{sat}$ is measured. We combine Equations (3) and (4) to arrive at a final rescaled form for $c_{sat}$ plotted against NCPR to assess the extent to which the data can be collapsed onto a master V-shaped profile. The final rescaled form of $c_{sat}$ takes the form:

$$c'''_{sat} = c_{sat}(T_0)\exp\left[-\left(\frac{T - T_0}{m}\right)\right]\left[n_a^2\left(\lambda_{aa} - 3\lambda_K n_K\right) + n_a n_R\left(2\lambda_{aR} - 6\lambda_K n_K\right)\right]; \tag{5}$$

We applied Equation (5) to analyze the totality of variant-specific temperature dependent data for $c_{sat}$. The results are shown in Fig. S2e. Here, the dashed red lines show linear fits of each arm of the V-shaped plot. The associated Pearson $r$-values that quantify the linear correlation are also shown on the plot. In calculating the fits, the rescaled $c_{sat}$ values for a given variant are averaged to one value so that each variant is weighted equally. The $\lambda$-values in Equations (2) – (5) were found by optimizing the Pearson correlation coefficients of the linear fits while keeping $\lambda_{aa}$ fixed at unity. Accordingly, the fit of Equation (5) to all of the data has three free parameters *viz*., $\lambda_{aR}$, $\lambda_K$, and $m$. The parameters we obtain from regression analysis are: $\lambda_{aR} = 1.69$, $\lambda_K = 0.0479$, and $m = 8.26$ K.

We tested the accuracy of this model by overlaying rescaled $c_{sat}$ values, rescaled according to Equation (5), for variants that were not used in the optimization. The results are shown in Fig. 3a. We find that the magnitudes of the Pearson *r*-values that quantify the strengths of linear correlations are still at least 0.95. The key message that is uncovered from the analysis in Fig. 3a and Equation (5) is that it helps us unmask the sticker and spacer determinants of the driving forces for phase separation. Importantly, it helps identify the contributions of NCPR to the driving forces for phase separation of LCDs, even for LCDs that are not enriched in charged residues.

*Parameterization of the LASSI model using values for $\nu^{app}$ that are derived from fits of molecular form factors to SAXS data*: In our coarse-grained lattice-based simulations, we use a single bead per residue and the interactions are isotropic. Simulations for individual chain molecules were performed using LASSI at a single temperature (T = 50 in the dimensionless units used for the energies and temperatures). For each interaction energy matrix, we performed ten independent simulations per sequence construct, and each simulation includes $3\times10^9$ proposed Monte Carlo moves that combine the range of moves developed by Choi et al. [17].

The goal was to parameterize interaction energies such that the dimensions of single chains, calculated using simulated ensembles, yield values for $\nu^{app}$ that are in accord with the values inferred from fitting molecular form factors to measured SAXS data [5]. Distributions of inter-residue distances were used to estimate $\nu^{app}$ values based on the method of Meng et al. [18]. A Gaussian process Bayesian optimization method [19, 20] as adapted by Ruff et al. [21] was implemented to obtain interaction parameters. Here, we iteratively search parameter space for pairwise interaction energies such that ensembles yield values for $\nu^{app}$ that minimize the sum of the square residuals between those obtained from simulations and experiments (Fig. 5a). The parameter bounds used in the optimization were changed manually over the course of the optimization if and only if a given parameter was often close to the upper or lower bound. Over 200 iterations were performed. The final bounds on parameters and values are shown in Table S2. We note that for every parameter, the absolute difference between the final value and either of the bounds is at least 10% of the absolute final value, suggesting the bounds are adequately distanced.

**Table S2: Optimized pairwise interaction energies (see column 4) used in the computational stickers-and-spacers model.** The table shows the lower and upper bounds (in terms of magnitudes) that were placed on each of the interaction energies based on previous calibrations [1].

| Pairwise Interaction | Lower Bound | Upper Bound | Final Value |
|---|---|---|---|
| Y-Y | -20 | -35 | -22.7 |
| Y-F | -12 | -30 | -18.9 |
| F-F | -5 | -25 | -16.0 |
| R-Aro | -8 | -25 | -11.0 |
| X-X | -2 | -4 | -2.78 |

*LASSI simulations of LCD phase behavior*: Multi-chain LASSI simulations at various temperatures were performed to simulate phase separation and obtain quantitative coexistence curves. For each variant, 200 distinct chain molecules, each with 137 beads, were placed in a cubic lattice with length 120 lattice units. Previous calibrations have shown that the numbers of molecules used in each of the simulations are adequate to avoid problems due to finite size artifacts [17]. The initial volume fraction in each simulation was approximately 0.016. Coexisting dense and dilute phases are delineated by identifying the single largest cluster in the system, computing the volume fraction within this single largest cluster as a function of radial distance from the center of mass of the single largest cluster, and comparing the volume fractions inside and outside the single largest cluster to that of the bulk well-mixed, homogeneous system. The volume fractions of the coexisting dense and dilute phases are estimated by equalizing the chemical potentials across the interface that we obtain from the radial density profiles. All multi-chain simulations were performed in triplicate, and the calculated volume fractions were essentially identical across replicates. At low temperatures, the calculated dilute phase volume fractions tend to have more variability due to the decreased likelihood that chains are in the dilute phase.

To convert from simulation temperature and volume fraction to experimental temperature and volume fraction, fixed scaling factors of 5.6 and 0.6 were used, respectively, for all variants and simulations in this study. These scaling factors were chosen by comparing the simulation and experimental phase diagrams of the aromatic variants (Fig. 5c-d) and are in accord with the published approach of Martin et al. [1]. To convert volume fractions to mass concentrations, we assumed that a volume fraction of 1.0 corresponds to a mass concentration of 1310 mg/ml based on the approach of Wei et al. [22].

*Incorporating NCPR-based corrections into the LASSI simulations*: The binodals shown in Fig. 6a-c incorporate a mean-field NCPR-based term whereby the NCPR of the variant determines the extent to which interactions are uniformly strengthened or weakened by decreasing or increasing the effective temperature, respectively. Depending on the NCPR, the simulation temperature is rescaled to $T^*$ according to the equation: $T^* = T + (a\Delta_{\text{NCPR}} + b)$. Here, $T$ is mapped to a new value $T^*$ based on the value of $\Delta_{\text{NCPR}} = |q_{\text{variant}} - q_{\text{mid}}|$, where $q_{\text{variant}}$ is the NCPR of the variant of interest, whereas $q_{\text{mid}}$, $a$, and $b$ are constants. The mean-field correction term ($a\Delta_{\text{NCPR}} + b$) can be visualized as an absolute-value function whose minimum vertex is at ($q_{\text{mid}}$, $b$) and slope is $-a$ to the left of the minimum and $+a$ to the right of the minimum. The value of $q_{\text{mid}}$ corresponds to the NCPR of the +4D variant (0.0287) since the +4D variant has the minimum residual from the aromatic fit in Fig. 4d. The value of $a = 1.54$, and this is based on the slopes of the linear fits in Fig. 4d and the slopes of the dilute arms of the LASSI-derived coexistence curves in Fig. 5c-d. The value of $b = -0.045$ to ensure equivalence of the effective and actual temperatures. Specifically, the value of $b$ ensures that the mean-field correction is zero for the WT A1-LCD and the aromatic variants, where the NCPR does not change vis-à-vis the WT. The approach of incorporating NCPR-based corrections by rescaling the simulation temperatures provides a way to account for the destabilizing effects of ionizable residues without explicitly modeling them as unique, highly solvated spacers. Note that this approach only works for obtaining NCPR-based corrections to computed binodals. Descriptions of single-chain conformations in the dilute phase do not incorporate these corrections.

*Error analysis to quantify deviations between computed and measured binodals*: We evaluate the error of our LASSI-derived binodals by comparing to experimentally derived binodals using a multi-step process:

(1) Perform a linear regression analysis of $\log_{10}(c_{\text{sat}})$ versus temperature for LASSI-derived dilute arms.

(2) For each temperature $T$, we calculate $x(T) = \left( \frac{c_{\text{sat, LASSI}}(T)}{c_{\text{sat,Expt.}}(T)} \right)$. Here, $c_{\text{sat,LASSI}}$ and $c_{\text{sat,Expt.}}$ are values of $c_{\text{sat}}$, in molar units, that are derived from LASSI simulations, from step (1), and experiments, respectively.

(3) Calculate the exponential root mean square log (ERMSL) of $x$ as $\text{ERMSL} = \exp\left( \sqrt{\left\langle \left[ \ln x \right]^2 \right\rangle} \right)$.

Here, the angular bracket denotes an average over all values of $[\ln x]^2$. The ERMSL is a positive value greater than or equal to 1 and can be interpreted as a measure of the error factor between the simulation-derived and experiment-derived binodals, specifically the dilute arms. For example, an ERMSL value of 10 indicates that, on average, the $c_{\text{sat}}$ values differ by about a factor of 10, or one order of magnitude. Alternatively, an ERMSL value of 1 indicates that there is no error between the dilute arms and that they should overlay perfectly. We report the ERMSL of each variant in Fig. 5e and Fig. S12d. In Fig. S12d, we specifically report the ERMSL of each variant with and without incorporation of the mean-field NCPR-based term and find that after incorporation of the mean-field term, the highest ERMSL value is only about 3.5. The ERMSL is akin to the root mean square log error (RMSLE) often used in machine learning, except for two important distinctions: First, when calculating the RMSLE, 1 is added to both the numerator and the denominator of the argument ($x$ in our case). This is done to prevent an indeterminate value of ln(0). In our case, we do not need to include this bias since the $c_{\text{sat}}$ values are necessarily greater than zero. Second, unlike with RMSLE, we take the exponential of our final value to bring the error back to an interpretable scale. This exponential operator is a reciprocal function of the inner logarithmic operator, in the same way that the outer square root operator is a reciprocal function of the inner square operator. As for the dense arms of the calculated binodals, the data show good agreement with measured binodals. However, given the complexity of these measurements, we do not have experimental data across the full range of temperatures for all variants, and therefore a quantitative analysis of computed and measured right arms of binodals cannot be performed across all variants.

*Overlap concentration calculation*: We calculate the overlap concentration of simulated constructs using the approach of Wei et al. [22]. Specifically, we use the equation $\phi^* = \frac{Nr^3}{\left(\sqrt{\left\langle R_e^2 \right\rangle}\right)^3}$ where $N$ is the number of residues in each chain, $r$ is the radius of each residue, and $\left\langle R_e^2 \right\rangle$ is the mean-squared end-to-end distance of a single chain. Here, we set $N$ = 137, $r$ = 0.5 lattice units. We apply the equation for calculations of volume fractions using end-to-end distances from simulations of single chains.

**Table S3: Estimated values for $R_g$ and $\nu^{app}$ from analysis of SAXS data using an empirical molecular form factor.**

| Construct | $R_g$ (Å) | $R_g$ error | $\nu^{app}$ | Standard error in estimate of $\nu^{app}$ |
|---|---|---|---|---|
| A1-LCD$^{-NLS}$ | 27.60 | 0.16 | 0.442 | 0.006 |
| A1-LCD$^{+NLS}$ | 25.83 | 0.11 | 0.430 | 0.004 |
| A1-LCD$^{-12F+12Y}$ | 26.04 | 0.20 | 0.429 | 0.007 |
| A1-LCD$^{+7F-7Y}$ | 27.18 | 0.13 | 0.454 | 0.006 |
| A1-LCD$^{-9F+6Y}$ | 26.55 | 0.10 | 0.457 | 0.005 |
| A1-LCD$^{-8F+4Y}$ | 27.07 | 0.07 | 0.461 | 0.003 |
| A1-LCD$^{-9F+3Y}$ | 26.83 | 0.13 | 0.460 | 0.006 |
| A1-LCD$^{-10R}$ | 26.71 | 0.07 | 0.468 | 0.004 |
| A1-LCD$^{-6R}$ | 25.73 | 0.09 | 0.448 | 0.004 |
| A1-LCD$^{+2R}$ | 26.23 | 0.23 | 0.440 | 0.009 |
| A1-LCD$^{+7R}$ | 27.09 | 0.07 | 0.442 | 0.003 |
| A1-LCD$^{-3R+3K}$ | 26.34 | 0.15 | 0.447 | 0.006 |
| A1-LCD$^{-6R+6K}$ | 27.87 | 0.08 | 0.467 | 0.003 |
| A1-LCD$^{-10R+10K}$ | 28.49 | 0.05 | 0.480 | 0.002 |
| A1-LCD$^{-4D}$ | 26.42 | 0.12 | 0.446 | 0.005 |
| A1-LCD$^{+4D}$ | 27.18 | 0.30 | 0.453 | 0.013 |
| A1-LCD$^{+8D}$ | 26.85 | 0.07 | 0.437 | 0.003 |
| A1-LCD$^{+12D}$ | 28.01 | 0.12 | 0.451 | 0.004 |
| A1-LCD$^{+12E}$ | 28.52 | 0.05 | 0.457 | 0.002 |
| A1-LCD$^{+7K+12D}$ | 29.21 | 0.08 | 0.467 | 0.003 |
| A1-LCD$^{+7K+12D\ blocky}$ | 25.62 | 0.14 | 0.412 | 0.005 |
| A1-LCD$^{-12F+12Y-10R}$ | 26.07 | 0.20 | 0.466 | 0.011 |
| A1-LCD$^{-10F+7R+12D}$ | 28.60 | 0.04 | 0.466 | 0.002 |

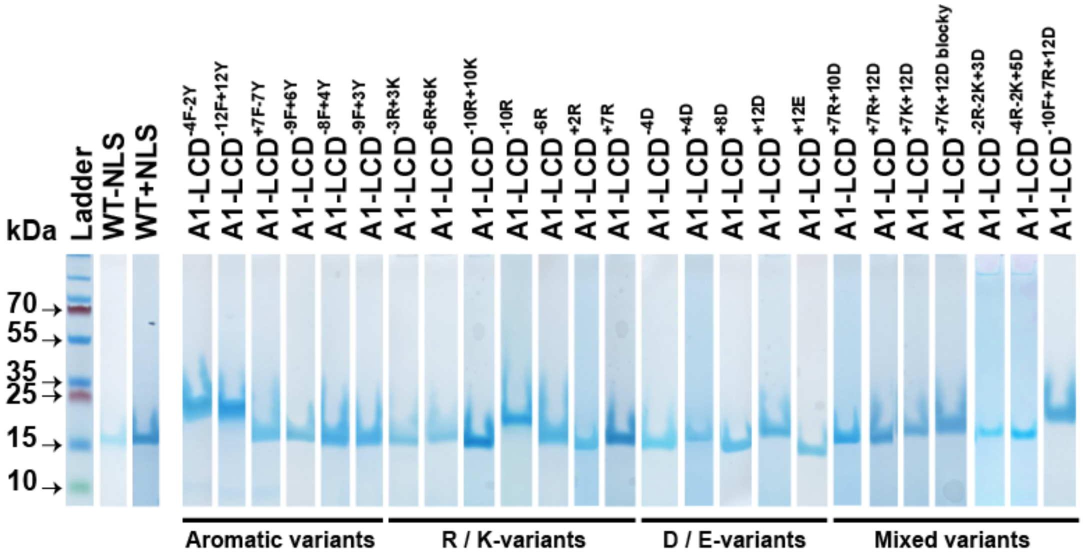


**Fig. S1: SDS-PAGE of purified A1-LCD variants used in this study.**

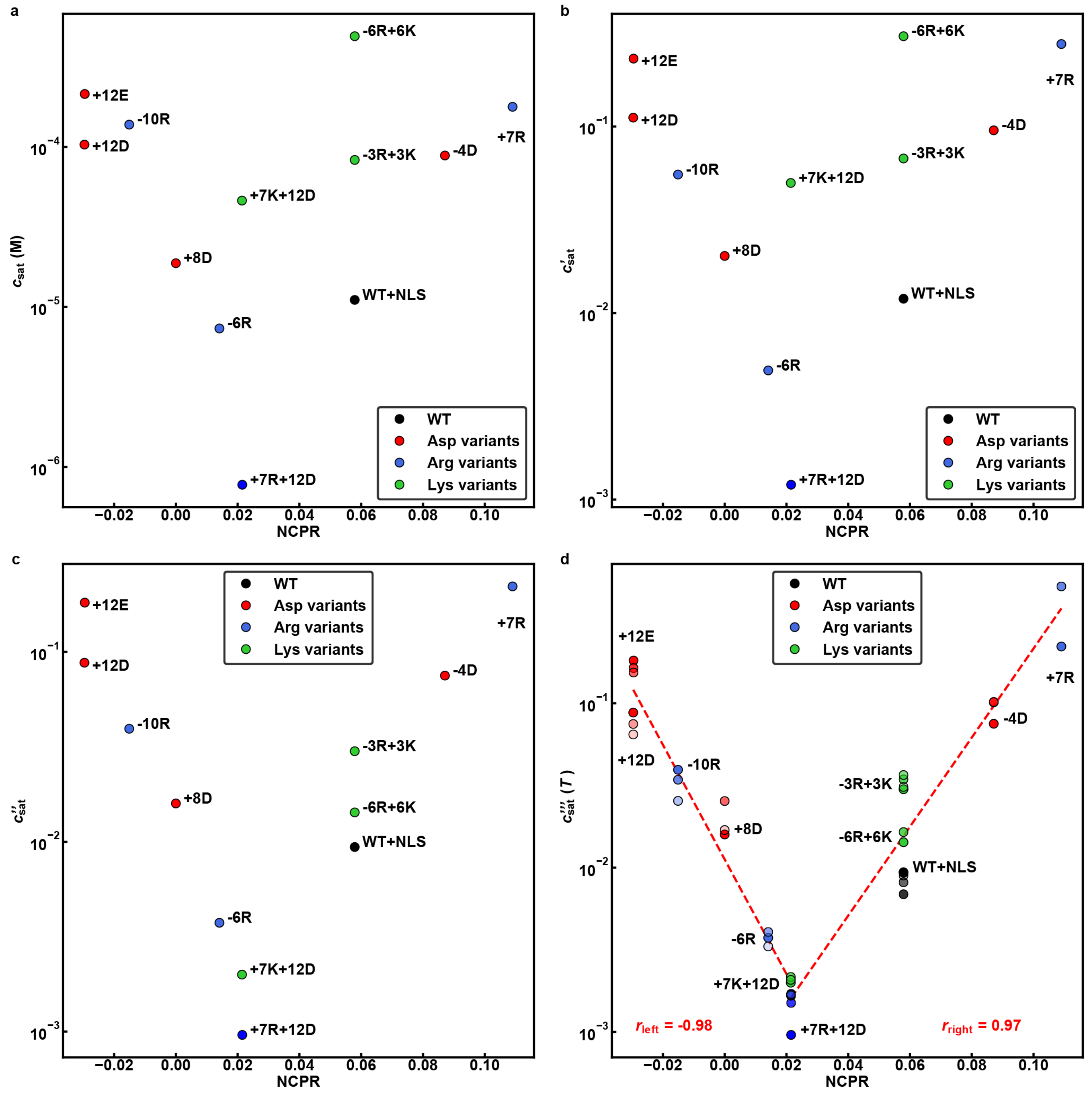


**Fig. S2: $\log_{10}(c_{sat})$ versus NCPR. (a)** Raw data for $\log_{10}(c_{sat})$ versus NCPR for all of the variants. **(b)** Results of rescaling $c_{sat}$ to account for the effects of Arg stickers. **(c)** Results of rescaling that account for the destabilizing effects of Lys in addition to accounting for the effects of Arg stickers. **(d)** Results of rescaling data using the approaches that lead to panels (b) and (c) combined with an accounting for the temperature-dependent variations to $c_{sat}$. The final result is a master curve onto which all $c_{sat}$ data for all measured temperatures collapse. The legends show the Pearson $r$-values that quantify the negative and positive linear correlations of the left and right arms, respectively.

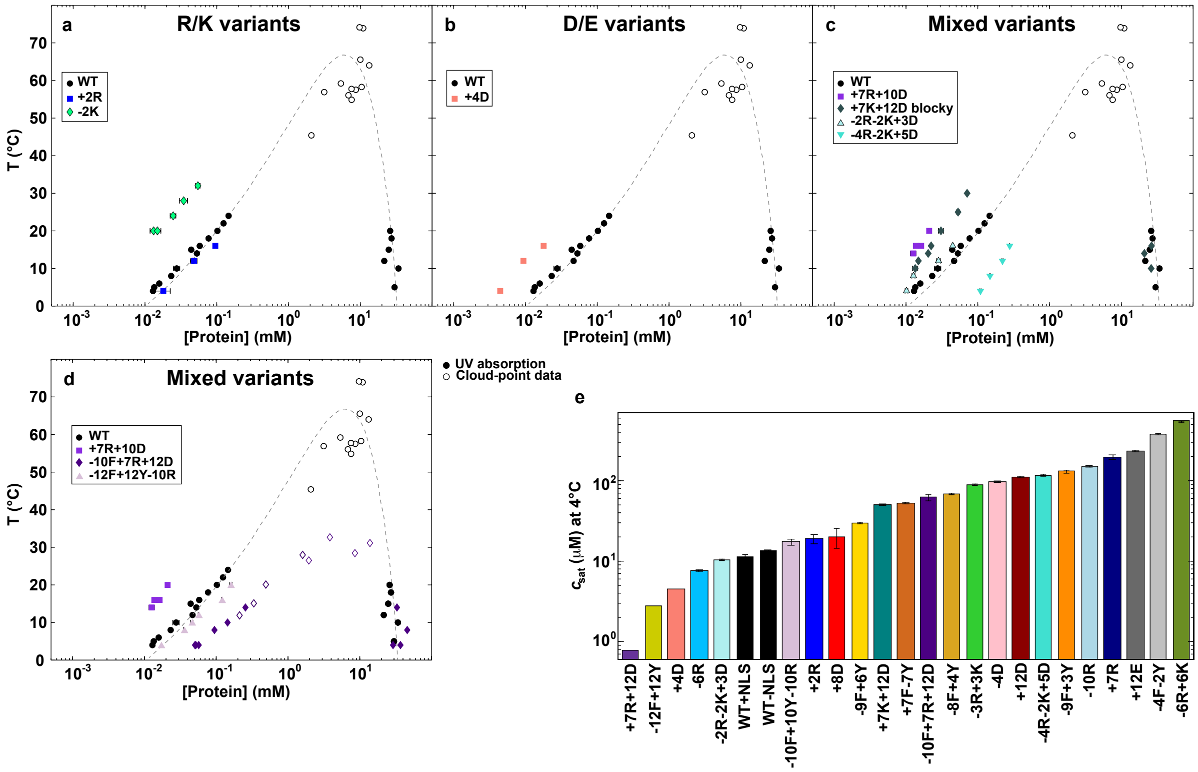


**Fig. S3: Measured binodals of additional A1-LCD variants designed to query the robustness of the master, V-shaped plot in Fig. 3a and the coarse-grained model in Fig. 6.** Binodals were measured using two different types of experiments, i.e., centrifugation followed by UV absorbance measurements (filled symbols) and cloud point measurements via static light scattering measurements (open symbols). The dashed line is a fit of Flory-Huggins theory to the experimental data for the WT A1-LCD to guide the eye.

- **(a)** Measured binodals for R/K variants that increase the positive NCPR.
- **(b)** Measured binodals for D/E variants that increase the negative NCPR.
- **(c)** Measured binodals mixed charge variants – details are in Fig. 2a.
- **(d)** Measured binodals of mixed charge variants to test coarse-grained model in Fig. 6.
- **(e)** Summary of all variant-specific $c_{sat}$ values measured at 4°C. Error bars indicate the standard error of replicate measurements.

a

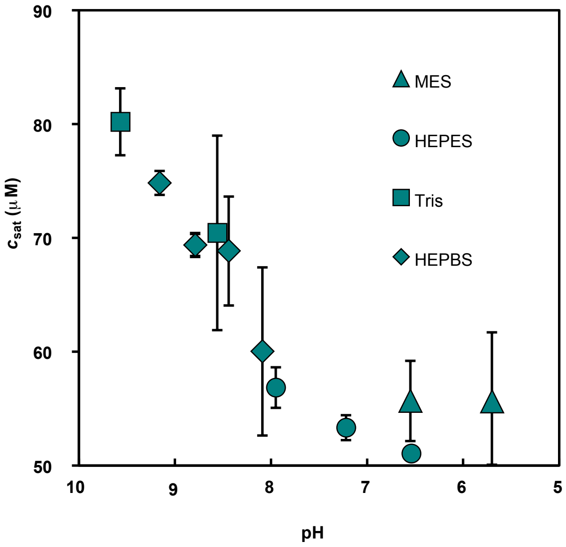


b

| Measured pH | Calculated charge | #Lys | Protonated Lys |
|---|---|---|---|
| 5.70 | 3.341 | 9 | 9 |
| 6.54 | 2.948 | 9 | 9 |
| 6.55 | 2.944 | 9 | 9 |
| 7.22 | 2.639 | 9 | 8.98 |
| 7.95 | 2.113 | 9 | 8.85 |
| 8.06 | 1.998 | 9 | 8.78 |
| 8.44 | 1.645 | 9 | 8.54 |
| 8.56 | 1.481 | 9 | 8.40 |
| 8.79 | 1.056 | 9 | 8.01 |
| 9.16 | -0.14 | 9 | 6.84 |
| 9.57 | -2.634 | 9 | 4.36 |

**Fig. S4: Measured pH-dependence of $c_{sat}$ validates the prediction that the minimum value for $c_{sat}$ is realized at positive values of NCPR.**

**(a)** Saturation concentration of +7K+12D was measured at 4°C as a function of pH.

**(b)** Table summarizing the theoretical net charge of +7K+12D and the number of Lys residues that are calculated to be protonated at each pH.

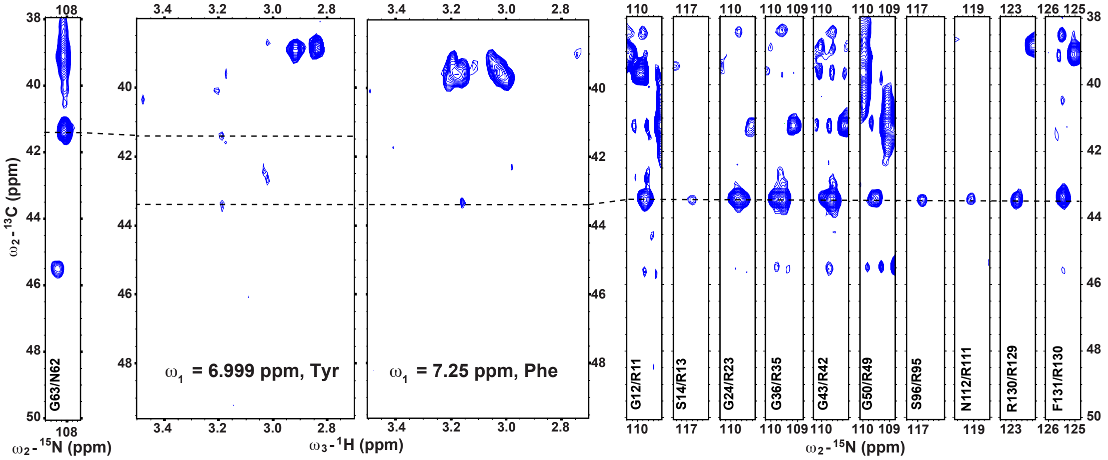


**Fig. S5: Additional NOEs involving aromatic residues in A1-LCD Δhexa.**

Tyr and Phe sidechain planes of $^{13}$C-resolved $^{1}H^{aromatic}$-$^{1}H^{aliphatic}$ NOESY (middle) show NOEs between sidechain protons of Tyr/Phe and Asn (left) and $H^{\delta}$ protons of Arg (right), respectively. A single (H)CC(CO)NH strip (corresponding to G63/N62) corresponds to the upfield carbon NOE resonance frequency. Degenerate Arg $C^{\delta}$ resonance frequencies (shown as strips of (H)CC(CO)NH spectrum on the right; see also Fig. 3e) match the downfield NOE frequency. The other NOEs could not be unambiguously assigned. NMR spectra of the dilute phase of A1-LCD Δhexa were recorded at 40°C.

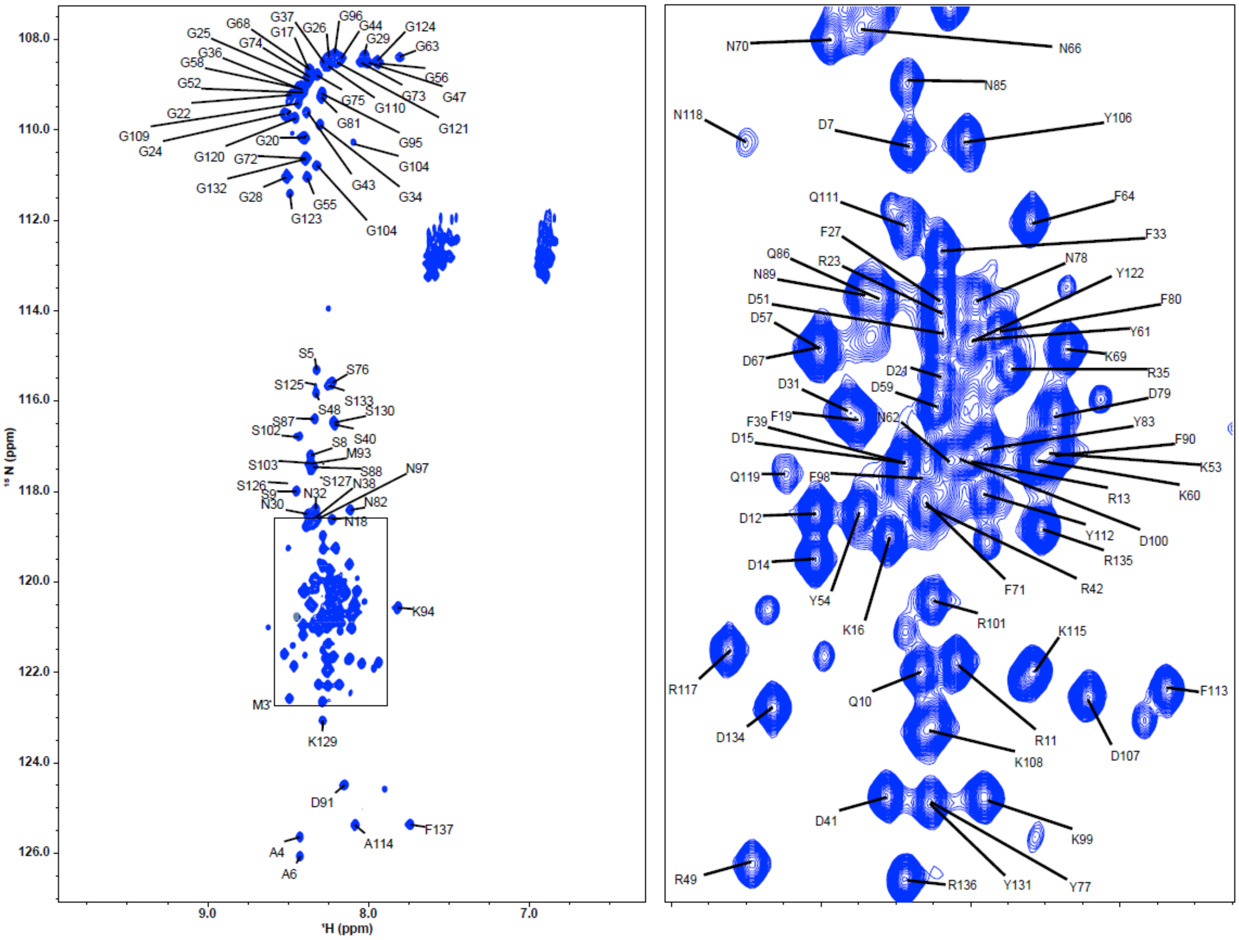


**Fig. S6: ¹H,¹⁵N HSQC spectrum of the A1-LCD variant +7K+12D.** The plot on the right is an expansion of a crowded area of the spectrum (indicated by a box on the left).

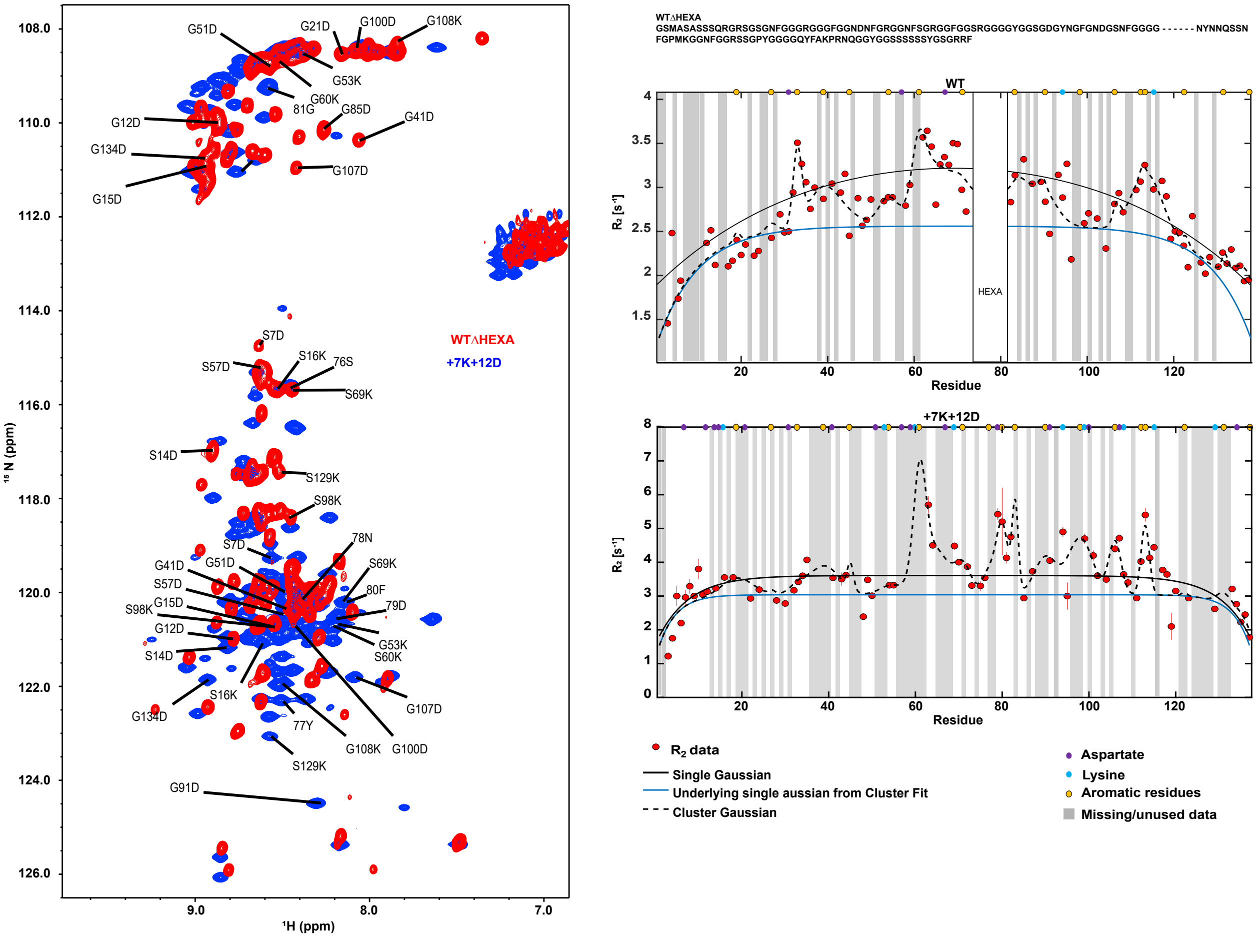


**Fig. S7: Aromatic residues are the main stickers in A1-LCD variant +7K+12D.**

Panel on the left shows an overlay of $^{1}H,^{15}N$ HSQC spectra of WT A1-LCD Δhexa variant (missing residues 259-264, red) as reported in (Martin et al. [1]) and +7K+12D (blue). The amino acid substitutions result in chemical shift changes across the spectrum. The panel on the right shows $^{15}N$ $R_2$ relaxation profiles for WT A1-LCD Δhexa (top) and the +7K+12D variant (bottom). The solid black profile represents a pure Gaussian-like fit, whereas the black dashed fit represents multiple regions of enhanced relaxation centered at aromatic residues (yellow) with the blue line representing the underlying Gaussian profile from this fit with a persistence length of 7.8 amino acid residues.

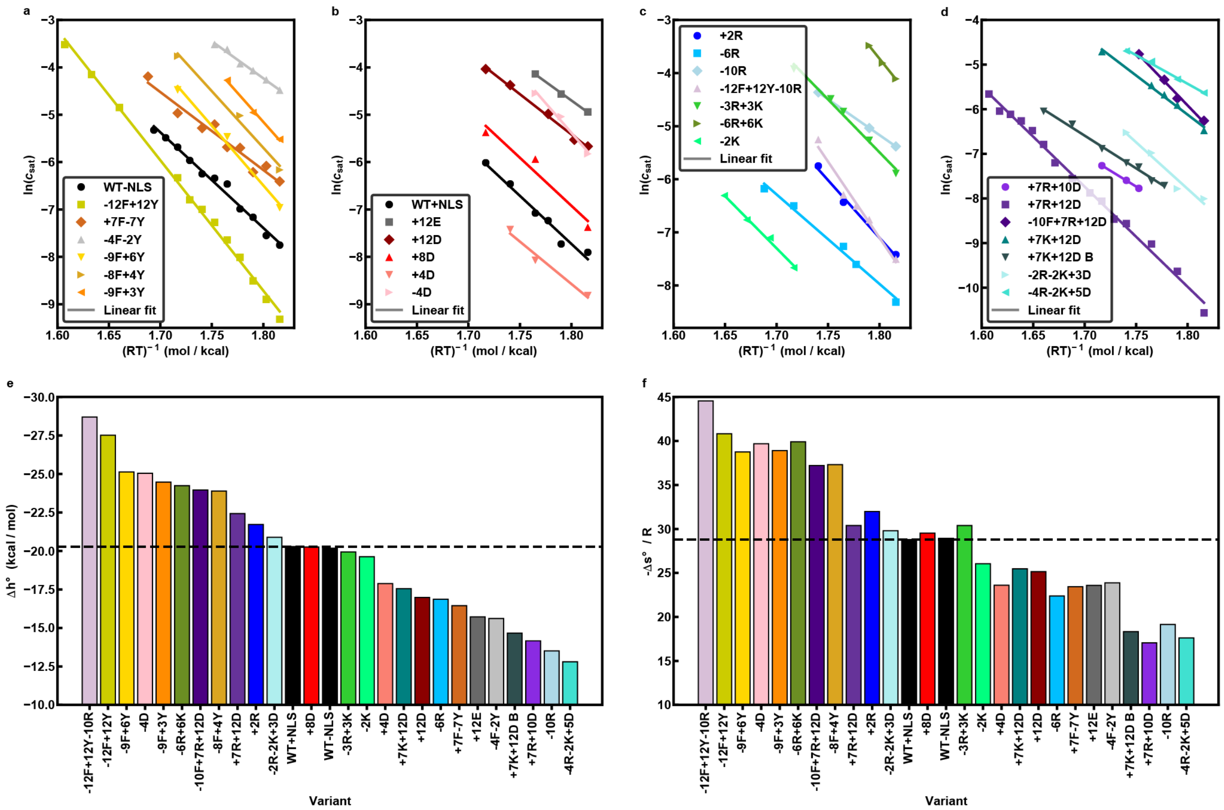

**Fig. S8: van't Hoff analysis for all of the A1-LCD variants**

Panels **(a-d)** show plots of ln($c_{sat}$) vs. $(RT)^{-1}$ for **(a)** aromatic variants, **(b)** Asp/Glu variants, **(c)** Arg/Lys variants, and **(d)** mixed charge variants. We do not include data for the +7R and -12F+12Y+7R variants because of the near critical behavior of these variants across the concentration and temperature ranges that we investigated. Although we use data for all of the variants, we stress that the analysis is reliable if and only if we have at least five different temperatures where $c_{sat}$ values are measured. Panels **(e)** and **(f)** show variant-specific estimates for $\Delta h^\circ$ and $-\Delta s^\circ/R$ extracted from panels (a) – (d) using the van't Hoff analysis. Here, $R = 1.98717\times10^{-3}$ kcal/mol*K.

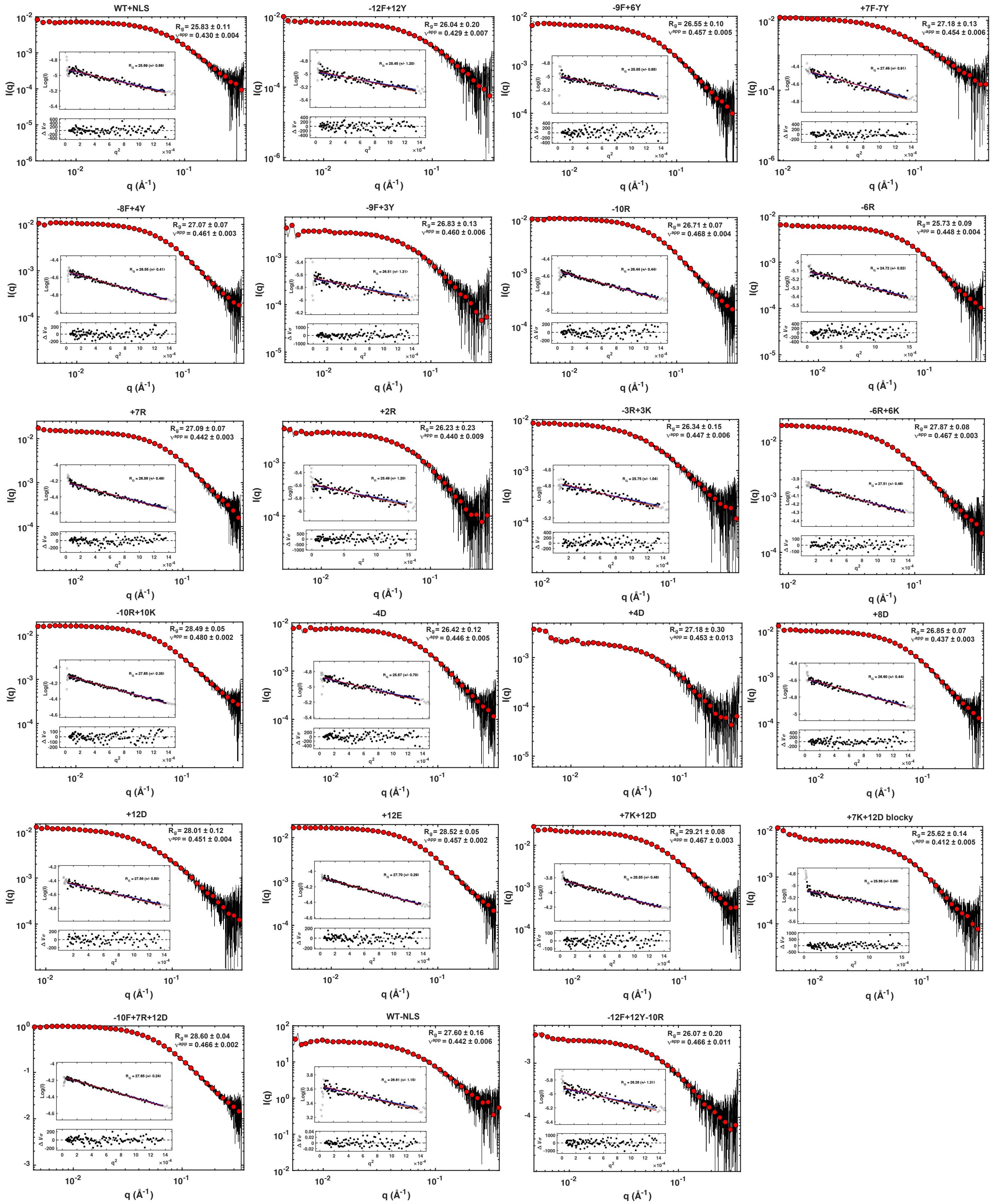


**Fig. S9: Raw SAXS data of all A1-LCD variants.**

SAXS data for all A1-LCD variants that were analyzed in this manner presented as I(q) versus q normalized by the forward scattering. The raw data (black) is overlaid with logarithmically smoothed data for visualization (red circles). The results from the fit to the empirical MFF [5] are indicated in the upper right corner. (Results are summarized in Table S3). The inset is the Guinier fit with the resulting $R_g$. Deviations from the linearity in the Guinier region prevented a Guinier fit for variant +4D; a fit to the MFF was possible, nonetheless.

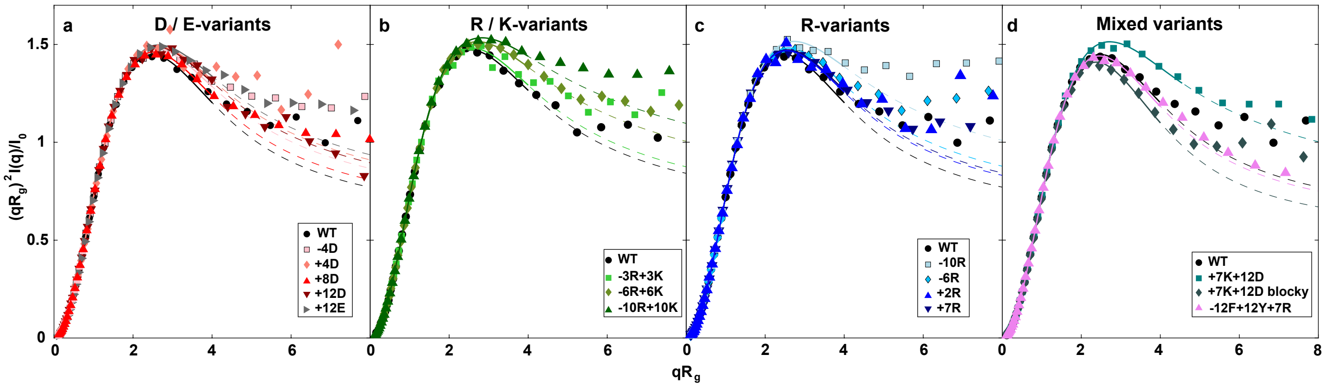


**Fig. S10: Kratky plot of the raw SEC-SAXS of A1-LCD variants.**

Data are shown for **(a)** D / E variants, **(b)** R / K variants, **(c)** R-variants, and **(d)** mixed variants. Data were logarithmically smoothed into 40 bins. Solid lines are fits to system-specific empirical molecular form factors (MFF).

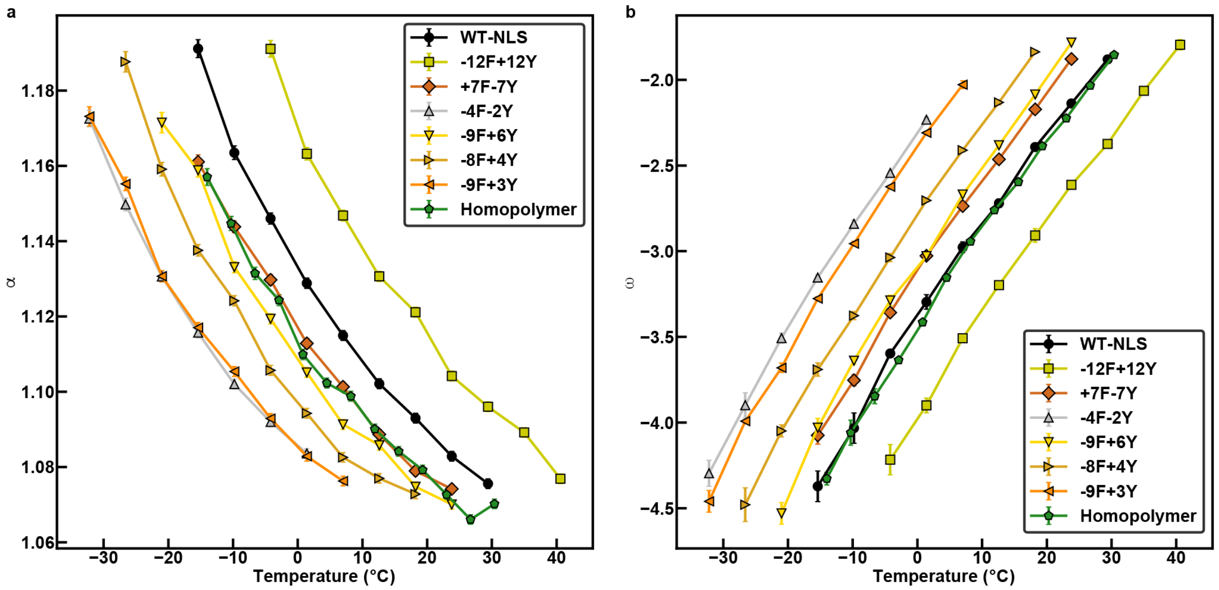


**Fig. S11: Analysis of swelling ratios and widths of two-phase regimes for aromatic variants.**

**(a)** Plots of the swelling ratio $\alpha$ against temperature for the WT A1-LCD (black), aromatic variants, and the equivalent homopolymer (green).

**(b)** Plots of the width of the two-phase regime $\omega$, defined as $\log_{10}$ of the ratio of the dilute phase concentration to the dense phase concentration. Here, we show data for $\omega$ plotted against temperature for the WT A1-LCD (black), aromatic variants, and the equivalent homopolymer (green).

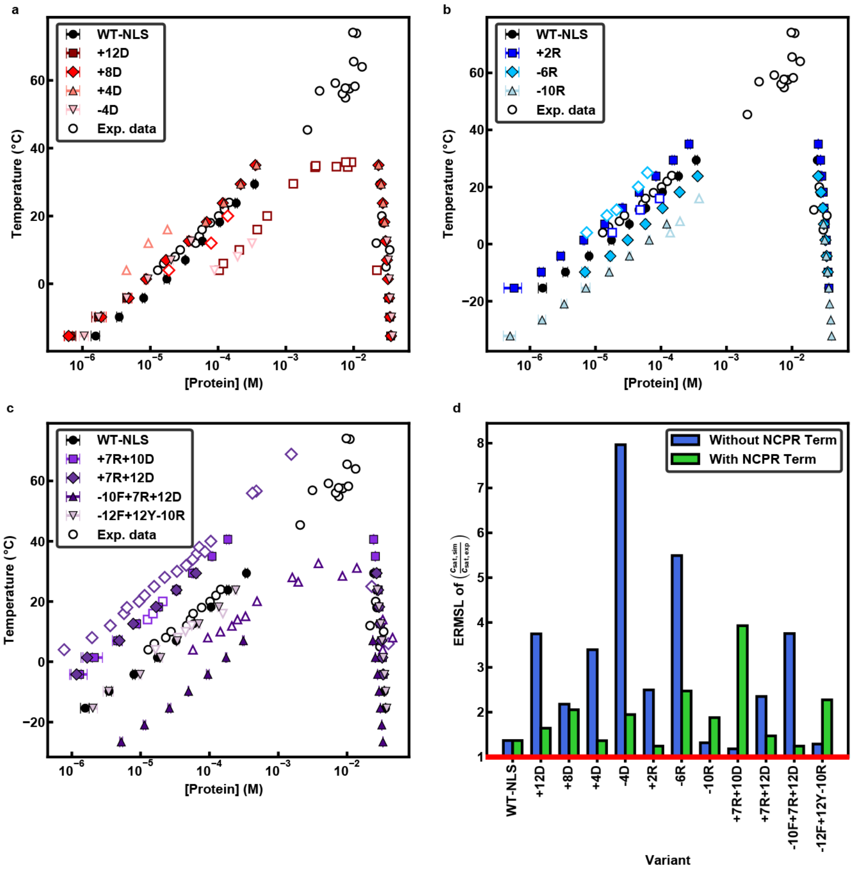


**Fig. S12: Comparisons of calculated and measured binodals for charge variants.** Results are shown for calculations that do not use a mean-field NCPR-based correction.

**(a-c)** Computed (solid circles) and measured (open circles) binodals of Asp variants (a), Arg variants (b), and variants not used in the model parameterization (c).

**(d)** Average factor of error of simulation-derived phase diagrams from experiment-derived phase diagrams of A1-LCD charge variants with and without the mean-field NCPR-based simulation parameter. A value of 1 indicates no error and a value of 10 indicates one magnitude of error. See methods for detailed explanation of exponential root mean square logarithm (ERMSL).

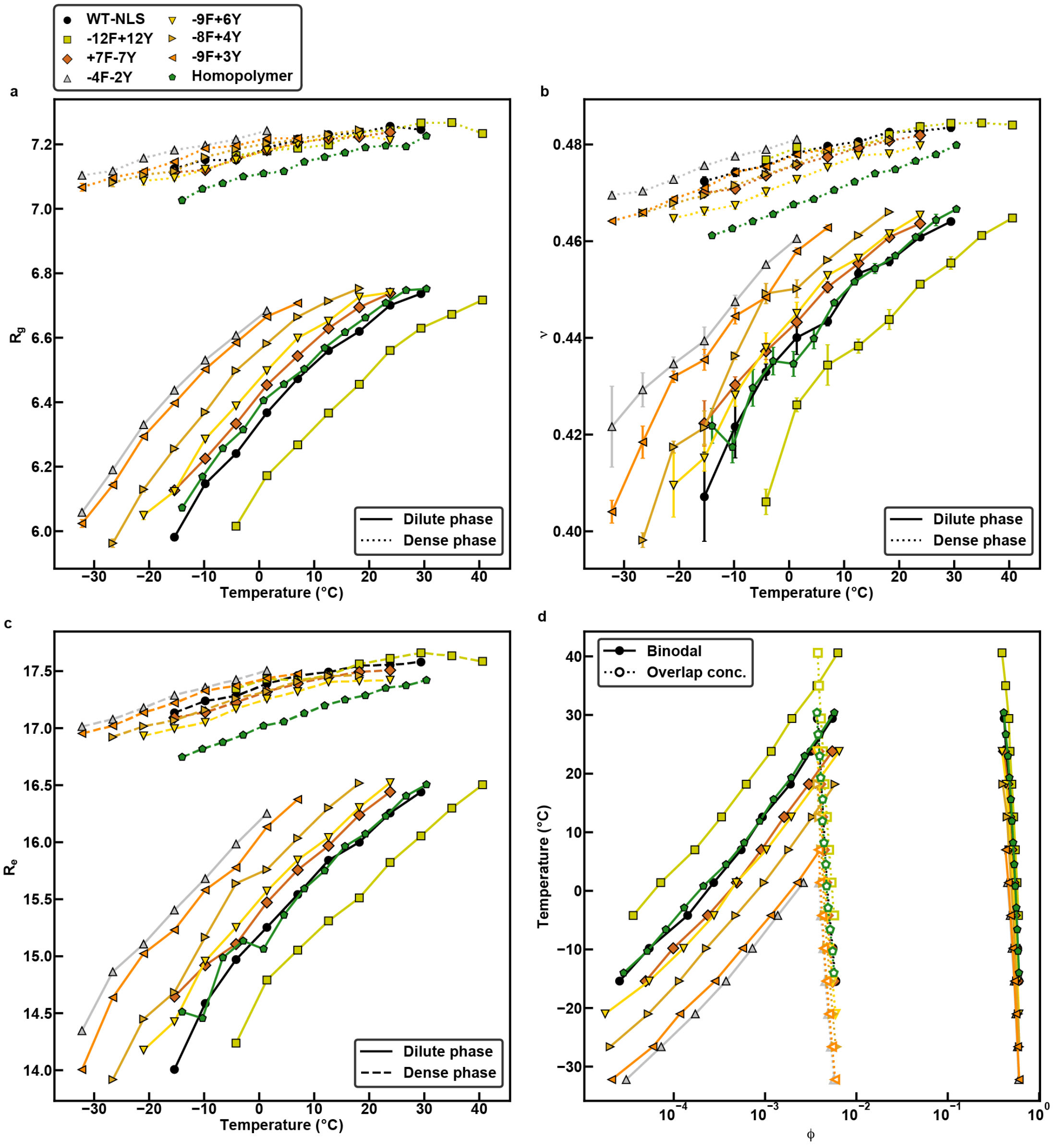


**Fig. S13: Dense phase and dilute phase comparisons from simulations of the aromatic variants and a homopolymer equivalent of WT A1-LCD.**

**(a)** Root-mean-square radii of gyration ($R_g$), calculated in lattice units, plotted against temperature for WT A1-LCD (black), aromatic variants, and the equivalent homopolymer (green) in the dilute (solid lines) and dense phases (dashed lines).

**(b)** Inferred average values for the apparent scaling exponent $\nu^{app}$ plotted against temperature for dimensions of single chains in the dilute (solid lines) and dense phases (dashed lines). Results are shown for WT A1-LCD (black), aromatic variants, and the equivalent homopolymer (green).

**(c)** Average end-to-end distance ($R_e$) of individual chain molecules in dilute (solid lines) and dense phases (dashed lines) plotted against temperature for WT A1-LCD (black), aromatic variants, and the equivalent homopolymer (green).

**(d)** Calculated binodals (solid markers; solid lines) and overlap concentrations (open markers; dashed lines) for WT A1-LCD (black), aromatic variants, and the equivalent homopolymer (green).

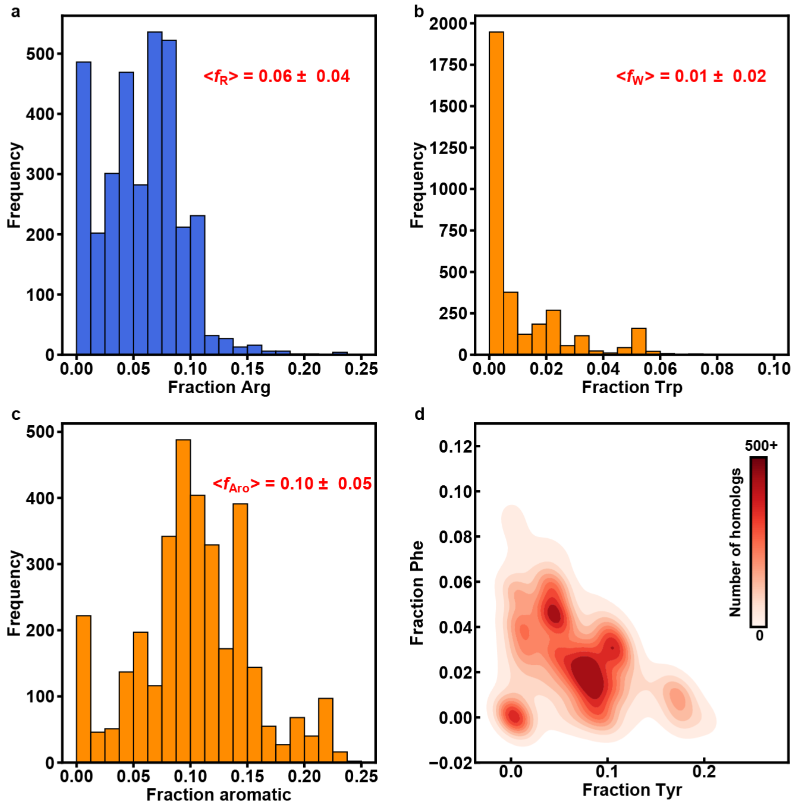


**Fig. S14: Compositional biases in LCDs drawn from homologs of the FUS / FET family of proteins.**

**(a)** Distribution of the fraction of Arg residues.

**(b)** Distribution of the fraction of Trp residues.

**(c)** Distribution of the and fraction of aromatic residues.

**(d)** Joint distribution of the fraction of Tyr and Phe residues.

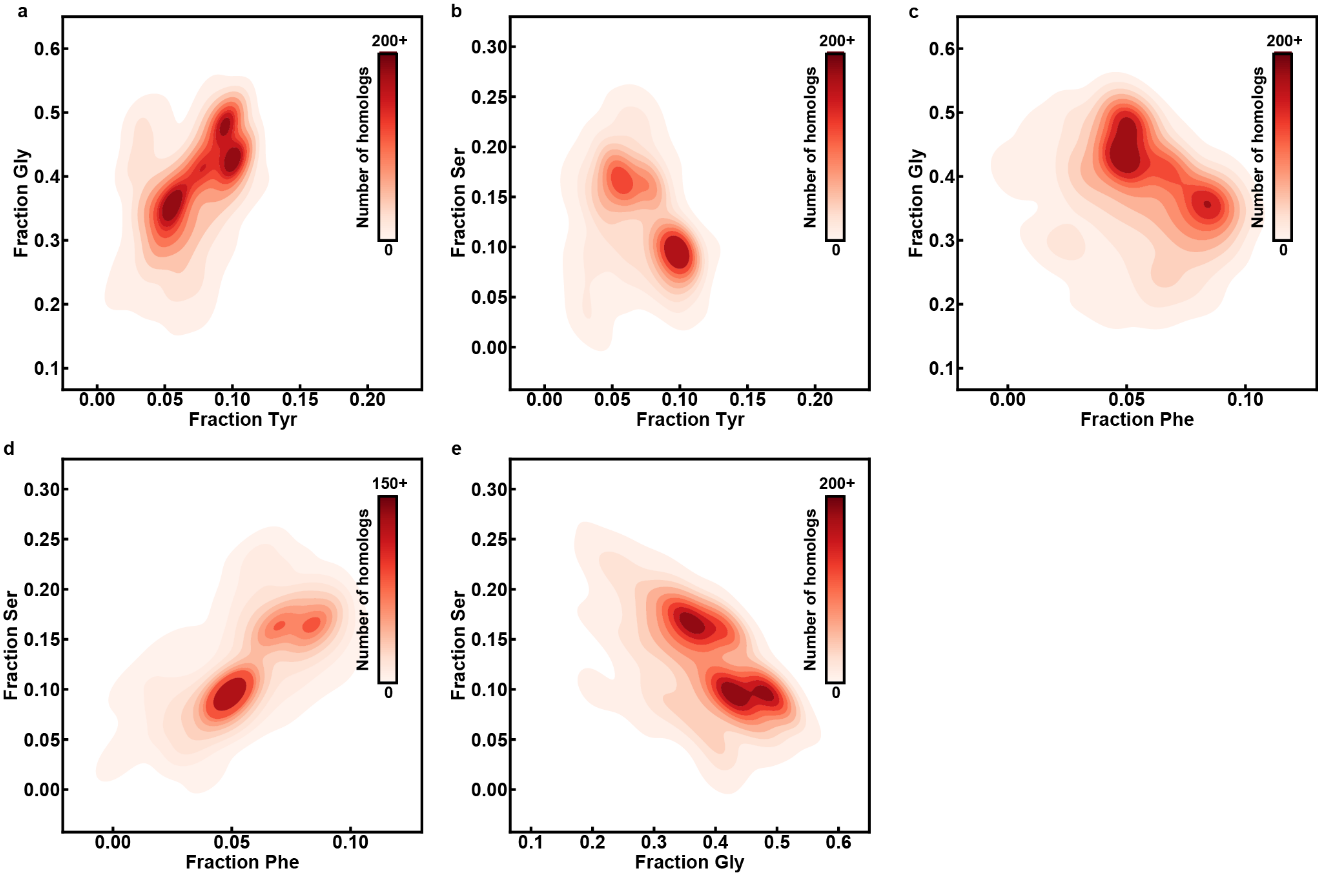


**Fig. S15: Covariation analysis of sticker and spacer contents across LCDs from 848 homologs of hnRNPA1**

(a) 2D histogram quantifying the joint distribution of the fractions of Tyr and Gly. This histogram shows a positive correlation between the fractions of Tyr and Gly.

(b) 2D histogram quantifying the joint distribution of the fractions of Tyr and Ser. This histogram shows a negative correlation between the fractions of Tyr and Ser.

(c) 2D histogram quantifying the joint distribution of the fractions of Phe and Gly. This histogram shows a negative correlation between the fractions of Phe and Gly.

(d) 2D histogram quantifying the joint distribution of the fractions of Phe and Ser. This histogram shows a positive correlation between the fractions of Phe and Ser.

(e) 2D histogram quantifying the joint distribution of the fractions of Gly and Ser. This histogram shows a negative correlation between the fractions of Gly and Ser.

References


1. Martin EW, Holehouse AS, Peran I, Farag M, Incicco JJ, Bremer A*, et al.* Valence and patterning of aromatic residues determine the phase behavior of prion-like domains. *Science* 2020, **367**(6478)**:** 694-699.

2. Martin EW, Hopkins JB, Mittag T. Small angle x-ray scattering experiments of monodisperse samples close to the solubility limit. *arXiv* 2020.

3. Kirby N, Cowieson N, Hawley AM, Mudie ST, McGillivray DJ, Kusel M*, et al.* Improved radiation dose efficiency in solution SAXS using a sheath flow sample environment. *Acta Crystallogr D Struct Biol* 2016, **72**(Pt 12)**:** 1254-1266.

4. Hopkins JB, Gillilan RE, Skou S. BioXTAS RAW: improvements to a free open-source program for small-angle X-ray scattering data reduction and analysis. *J Appl Crystallogr* 2017, **50**(Pt 5)**:** 1545-1553.

5. Riback JA, Bowman MA, Zmyslowski AM, Knoverek CR, Jumper JM, Hinshaw JR*, et al.* Innovative scattering analysis shows that hydrophobic disordered proteins are expanded in water. *Science* 2017, **358**(6360)**:** 238-241.

6. Delaglio F, Grzesiek S, Vuister GW, Zhu G, Pfeifer J, Bax A. Nmrpipe - a Multidimensional Spectral Processing System Based on Unix Pipes. *J Biomol Nmr* 1995, **6**(3)**:** 277-293.

7. Lee W, Tonelli M, Markley JL. NMRFAM-SPARKY: enhanced software for biomolecular NMR spectroscopy. *Bioinformatics* 2015, **31**(8)**:** 1325-1327.

8. Johnson BA. From Raw Data to Protein Backbone Chemical Shifts Using NMRFx Processing and NMRViewJ Analysis. *Methods Mol Biol* 2018, **1688:** 257-310.

9. Huerta-Cepas J, Szklarczyk D, Heller D, Hernández-Plaza A, Forslund SK, Cook H*, et al.* eggNOG 5.0: a hierarchical, functionally and phylogenetically annotated orthology resource based on 5090 organisms and 2502 viruses. *Nucleic Acids Research* 2018, **47**(D1)**:** D309-D314.

10. Sievers F, Wilm A, Dineen D, Gibson TJ, Karplus K, Li W*, et al.* Fast, scalable generation of high-quality protein multiple sequence alignments using Clustal Omega. *Molecular Systems Biology* 2011, **7**(1)**:** 539.

11. Consortium TU. UniProt: a worldwide hub of protein knowledge. *Nucleic Acids Research* 2018, **47**(D1)**:** D506-D515.

12. Holehouse AS, Das RK, Ahad JN, Richardson MOG, Pappu RV. CIDER: Resources to Analyze Sequence-Ensemble Relationships of Intrinsically Disordered Proteins. *Biophysical Journal* 2017, **112**(1)**:** 16-21.

13. Wang J, Choi JM, Holehouse AS, Lee HO, Zhang X, Jahnel M*, et al.* A Molecular Grammar Governing the Driving Forces for Phase Separation of Prion-like RNA Binding Proteins. *Cell* 2018, **174**(3)**:** 688-699 e616.

14. Semenov AN, Rubinstein M. Thermoreversible Gelation in Solutions of Associative Polymers. 1. Statics. *Macromolecules* 1998, **31**(4)**:** 1373-1385.

15. Choi J-M, Hyman AA, Pappu RV. Generalized models for bond percolation transitions of associative polymers. *Physical Review E* 2020, **102:** 042403.

16. Mahadevi AS, Sastry GN. Cooperativity in Noncovalent Interactions. *Chemical Reviews* 2016, **116**(5)**:** 2775-2825.

17. Choi J-M, Dar F, Pappu RV. LASSI: A lattice model for simulating phase transitions of multivalent proteins. *PLoS computational biology* 2019, **15**(10).

18. Meng W, Lyle N, Luan B, Raleigh DP, Pappu RV. Experiments and simulations show how long-range contacts can form in expanded unfolded proteins with negligible secondary structure. *Proceedings of the National Academy of Sciences* 2013, **110**(6)**:** 2123.

19. Rasmussen CE, Williams CKI. *Gaussian Processes for Machine Learning*. MIT Press: Cambridge, MA, 2006.

20. Pedregosa F, Varoquaux G, Gramfort A, Michel V, Thirion B, Grisel O*, et al.* Scikit-learn: Machine Learning in Python. *J Mach Learn Res* 2011, **12**(null)**:** 2825–2830.

21. Ruff KM, Harmon TS, Pappu RV. CAMELOT: A machine learning approach for coarse-grained simulations of aggregation of block-copolymeric protein sequences. *The Journal of Chemical Physics* 2015, **143**(24)**:** 243123.

22. Wei MT, Elbaum-Garfinkle S, Holehouse AS, Chen CC, Feric M, Arnold CB*, et al.* Phase behaviour of disordered proteins underlying low density and high permeability of liquid organelles. *Nature Chemistry* 2017, **9**(11)**:** 1118-1125.
